# Regulation of CHK1 inhibitor resistance by a c-Rel and USP1 dependent pathway

**DOI:** 10.1101/2021.05.26.445425

**Authors:** Jill E. Hunter, Amy E. Campbell, Nicola L. Hannaway, Scott Kerridge, Saimir Luli, Jacqueline A. Butterworth, Helene Sellier, Reshmi Mukherjee, Huw D. Thomas, Philip J. Brownridge, Johnathon Coxhead, Leigh Taylor, Peter Leary, Megan S.R. Hasoon, Ian Collins, Michelle D. Garrett, Claire E. Eyers, Neil D. Perkins

## Abstract

We previously discovered that deletion of c-Rel in the Eμ-Myc mouse model of lymphoma results in earlier onset of disease, a finding that contrasted with the expected function of this NF-κB subunit in B-cell malignancies. Here we report that c-rel -/- Eµ-Myc cells have an unexpected and major defect in the CHK1 pathway, with almost undetectable levels of CHK1 and CLSPN protein leading to therapeutic resistance to the highly specific CHK1 inhibitor (CHK1i) CCT244747. Similar downregulation of CHK1 levels was also seen in CCT244747 resistant U20S osteosarcoma cells. Further investigation revealed that downregulation of the deubiquitinase USP1 is responsible, at least in part, for these effects. Importantly, we demonstrate that c-rel -/- Eµ-Myc lymphoma cells survive though upregulation of compensatory PI3K/AKT pathway activity. Moreover, targeting this pathway with Pictilisib (GDC-0941) effectively killed c-rel -/- Eµ-Myc *in vivo,* while having no effect on wild type Eμ-Myc cells. This data reveals an NF-κB regulated pathway controlling CHK1 activity in cancer cells and identifies a potential mechanism for both acquiring and overcoming CHK1i resistance in cancer patients.

## Introduction

The development of targeted therapies has changed the face of cancer therapeutics, and allowed for a more personalised approach to treatment based on the molecular basis of an individual’s tumour. One of the oncogenic kinases identified as a ’druggable’ target is the checkpoint kinase, CHK1, and for this reason, CHK1 inhibitors (CHK1i) represent a potential new class of anti-cancer therapies, and are currently in clinical trials [1].

CHK1 plays a critical role in the response to DNA replication stress, which results from stalled DNA replication forks. In cancer cells, replication stress drives both genomic instability and clonal evolution [2–4]. It can be induced by a variety of mechanisms, including DNA damaging agents and by oncogenes such as MYC driving hyper-DNA replication [2–4]. Critical regulators of the cellular response to DNA replication stress, not only include CHK1 but also the kinase Ataxia Telangiectasia and Rad3 Related, which protect against tumorigenesis through promoting DNA repair [4, 5]. However, once established, tumour cells can also become addicted to this pathway since it enables them to survive on-going, potentially lethal, genomic instability. Therefore, inhibiting key protein kinases, such as CHK1, provides a potential therapeutic strategy in order to specifically target tumours that have become dependent on their activity [6].

We, and others have shown that there is significant crosstalk, between the ATR-CHK1 and the NF-κB pathways. For example, we have shown *in vitro* that CHK1 can phosphorylate the RelA subunit in its transactivation domain at Thr 505 (T505), resulting in inhibition of tumour promoting activities of NF-KB, including resistance to apoptosis, autophagy, cell proliferation and cell migration [7–11]. Direct phosphorylation of the p50 NF-KB subunit on Ser 329 by CHK1 was demonstrated following DNA damage [12, 13], and *in vitro* phosphorylation of p50 by CHK1 also regulates DNA binding of the p50 homodimer through phosphorylation on Ser 242, and homodimerisation through a phosphorylation event on Ser 337 [14]. Moreover, Kenneth et al [15], found that the c-Rel NF-KB subunit controls the expression of Claspin in cancer cell lines. This is of particular relevance as Claspin is an adaptor protein associated with DNA replication forks that is required for ATR dependent phosphorylation of CHK1 following DNA replication stress [16, 17].

Resistance to kinase inhibitors is a problem that is being frequently encountered in the clinic. Resistance can be either *de novo* or acquired, and there may be multiple different mechanisms that contribute to the resistance to particular agents. In the case of *de novo* resistance, this is primarily due to tumour intrinsic factors or patient specific pharmacokinetic factors such as the absorption, distribution, metabolism and excretion of the drug (reviewed in [18]).

Acquired resistance can develop *via* a variety of different mechanisms, such as the mutation of the drug target or a gene within a closely related signalling pathway. Histological changes, such as epithelial to mesenchymal transition [19] and increased growth factor production have also been reported to result in acquired resistance [20, 21]. However, one of the biggest contributing factors to acquired resistance is activation of compensatory signalling pathways, so that the cell can bypass the need for the inhibited target, often referred to as “bypass signalling” [18]. There are multiple strategies being successfully employed in the clinic, including the targeting of these compensatory signaling pathways, either alone or in combination with the initial targeted agent. This strategy has been used in mutant melanoma, where a combination of RAF and MEK inhibitors have shown efficacy [22].

Using an integrated ’omics-based approach we have investigated how *de novo* resistance to CHK1i is acquired and can be overcome, revealing that this is a two-step process. Initial downregulation of the CHK1 signalling pathway can occur by multiple different mechanisms; here we show that loss or inactivation of the CHK1 pathway either *via* loss of Claspin, or of the CHK1 deubquitinase (DUB), USP1, contributes to the downregulation of CHK1 signalling in our cell line and animal models. Secondly, we reveal upregulation of a compensatory PI3K/AKT signalling pathway in mice that are resistant to CHK1 inhibition which serves to drive continued tumour growth. Targeting this compensatory PI3K/AKT signaling pathway represents a potential therapeutic opportunity in patients developing resistance to CHK1i in the clinic.

## Results

### Eµ-Myc*/c-Rel^-/-^* lymphoma cells are resistant to CHK1 inhibition

Since over-expression of c-Myc is a feature of many types of cancer and results in DNA replication stress leading to genomic instability and tumorigenesis we chose the well-established Eµ-Myc mouse model of B-cell lymphoma [23] to investigate CHK1 inhibitor resistance mechanisms. We have previously found that knockout of the c-Rel NF-KB subunit in the Eµ-Myc model results in earlier onset of cancer [24]. We have also previously shown that the CHK1 inhibitor (CHK1i) SRA737, having just completed Phase I clinical trials (https://clinicaltrials.gov/ct2/show/NCT02797964), inhibits the growth of re-implanted wild type (WT) Eµ-Myc cells [25]. Since c-Rel has been described as an indirect regulator of CHK1 activity through inducing CLSPN gene expression [15], we were therefore curious as to whether deleting c-Rel would affect the response of Eµ-Myc lymphomas to CHK1 inhibition. We hypothesised that less efficient activation of CHK1 by ATR in response to MYC-induced DNA replication stress in c-rel -/- Eµ-Myc lymphoma cells could affect CHK1 inhibitor sensitivity.

We therefore evaluated the effectiveness of the CHK1i CCT244747 (the precursor compound to SRA737) *in vivo* by analysing its effect on the growth of five transplanted WT Eµ-Myc and Eµ-Myc/*c-Re/^-/-^* tumours. Each tumour, which originated from an independently derived spontaneous tumour bearing mouse, was implanted *via* the lateral tail vein into six syngeneic C57Bl/6 recipient mice and three were treated orally with CCT244747 once a day for nine days, while three received a vehicle control (Fig. 1A & S1A). Treatment commenced at the point at which tumours in the lymphoid organs became palpable. After treatment, we observed a striking reduction in lymphoid tumour burden in all mice re-implanted with WT Eµ-Myc lymphomas (Figs. 1B & C, S1B). By contrast, four of the five Eµ-Myc*/c-Re/^-/-^* lymphomas showed no significant reduction in lymphoid tumour burden after CCT244747 treatment, with one lymphoma only exhibiting a partial response in the thymus and cervical lymph nodes. The resistance of Eµ-Myc*/c-Re/^-/-^* lymphomas was confirmed *ex vivo*. Treatment of Eµ-Myc lymphoma cells with CCT244747 for 96 hours resulted in small but significant differences, with WT cells having reduced viability relative to Eµ-Myc*/c-Re/^-/-^* or Eµ-Myc/*Re/a*^T505A^ tumour cells (Fig. S1C). The reduced magnitude of the effects of CCT244747 seen here likely reflects the low level of proliferation seen with Eµ-Myc cells when cultured *ex vivo*. These data confirmed that regulation of CHK1/DNA replication stress by the c-Rel NF-κB subunit *in vivo* significantly affects the sensitivity of Eµ-Myc lymphoma cells to CHK1 inhibition.

**Figure 1.**
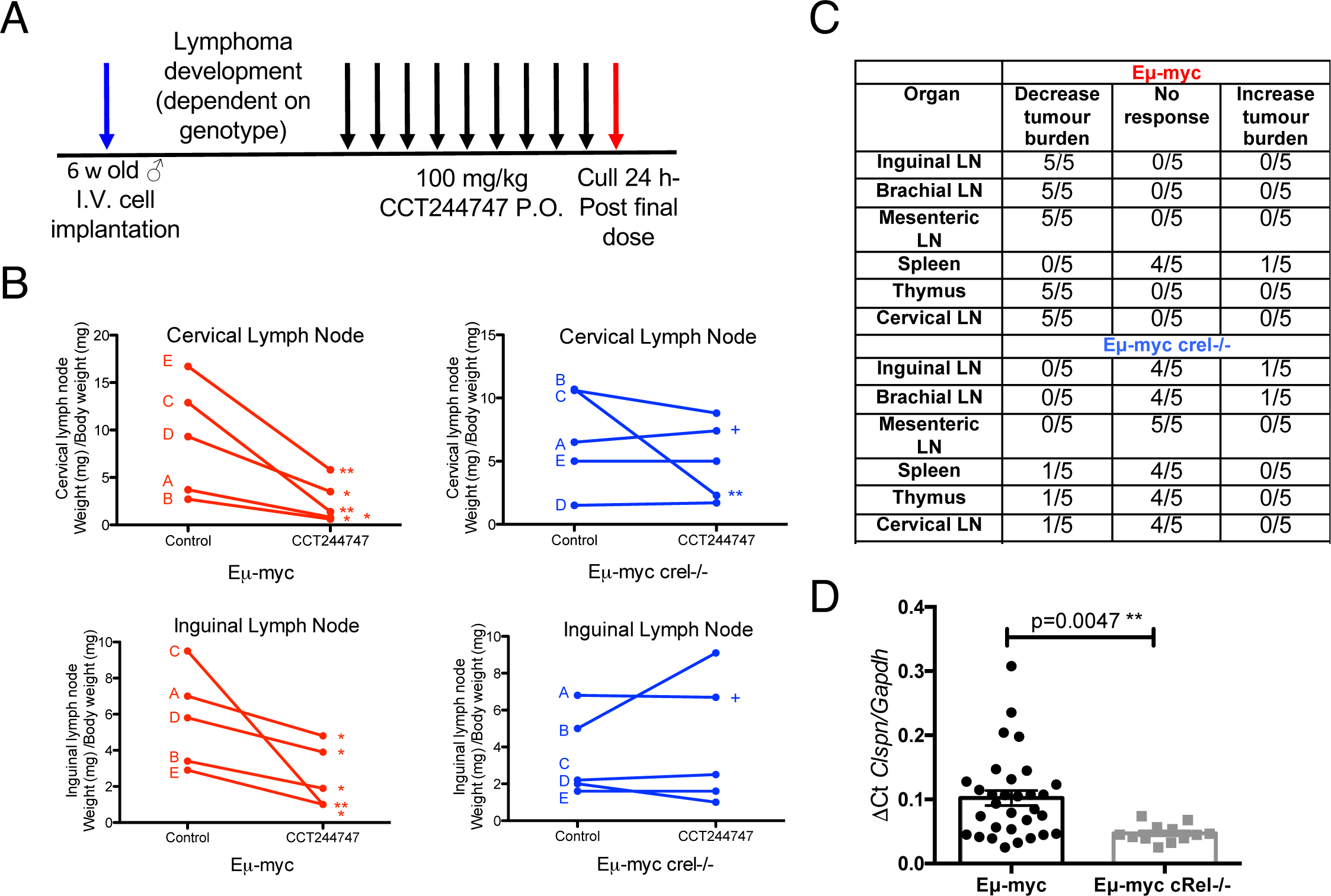
Claspin expression is reduced in Eµ-Myc*/c-rel-/-* and renders Eµ-Myc*/c-rel-/-* lymphoma resistant to Chk1 inhibition. (A) Schematic diagram illustrating the CHK1i *in vivo* study in Eµ-Myc and Eµ-Myc/*c-re/-/-* mice. 6 week old C57Bl/6 WT mice were implanted with either Eµ-Myc or Eµ-Myc/*c-re/-/-* (blue arrow) and once tumours became palpable were treated with either 100 mg/kg CCT244747 p.o or vehicle control once daily for 9 days (black arrows). Mice were euthanised 24 hours after the final dose (red arrow) and tumour burden assessed. (B) Line graphs showing the mean response of the five reimplanted Eµ-Myc [64] and Eµ-Myc crel-/- (blue) tumours and their response to CCT244747. Each of the 5 spontaneously derived tumours was implanted into 6 syngeneic recipient C57Bl/6 mice, 3 were treated with CCT244747 (100 mg/kg p.o), and 3 with vehicle control, for 9 days once lymphoid tumours became palpable. A response was defined as a significant change in tumour burden (P<0.05) using unpaired Student’s t-tests. The complete data set is summarised in (D). ’+’ indicates one experiment where treatment was stopped after 7 days and the mice were sacrificed early due to the mice becoming too ill. (C) Table showing the response of 5 re-implanted Eµ-Myc and Eµ-Myc/*c-re/-/-* tumours to CCT244747, in all sites where lymphoid tumour burden is anticipated in this model. (D) Q-PCR data showing relative Claspin expression in tumorigenic spleens from Eµ-Myc (n=30) and Eµ-Myc/*c-re/-/-* (n=11) mice. Data represents mean ± SEM. p** <0.01 (Unpaired Student’s t-test). Claspin expression is reduced in c-Rel^-/-^ tumours. Data represents mean ± SEM, each point is an individual mouse.

### Eµ-Myc lymphomas lacking c-Rel exhibit altered cell signalling and response to CHK1 inhibition

Since CLSPN can be an NF-κB target gene [15] we next investigated its expression and found that CLPSN mRNA expression was significantly down regulated in Eµ-Myc lymphoma cells lacking c-Rel (Fig 1D). By contrast, we did not observe any effect of deleting c-Rel on CLSPN expression in normal B-cells (Fig S1D). This loss of CLSPN expression in c-Rel^-/-^ Eµ-Myc lymphomas implied that ATR/CHK1 signalling might be compromised in these cells. We therefore decided to explore how these cells respond at an early time point to a single dose of CCT244747 *in vivo*. By examining this acute response, we reasoned that we could gain insights into how signalling in these cells had been rewired, something not possible with longer CCT244747 treatment where the mixture of dead, dying and surviving lymphoma cells was likely to confound analysis.

We therefore investigated the nature of the response of re-implanted WT and c-Rel^-/-^ Eµ-Myc lymphomas following acute treatment with the CHK1i, CCT244747, using a combination of (phospho)proteomic and RNA Seq analysis. To determine the optimum timepoint to study these changes, mice re-implanted with WT Eµ-Myc lymphomas were culled at various timepoints following a single dose of CCT244747 (Fig S2A). We observed a significant DNA damage response, as measured by an increase in phosphorylation of H2AX at Ser 139, after 8 hours of treatment that diminished at 24 and 48 hours (Fig 2A). Moreover, we observed high levels of CHK1 phosphorylation at Ser 345 at the 8-hour time point, an ATR-mediated phosphorylation site that typically becomes hyperphosphorylated upon CHK1 inhibition [26] 8 hours after treatment that diminished at 24 and 48 hours (Fig 2A). Hence we used this 8 hour timepoint in all further studies (Fig 2A).

**Figure 2.**
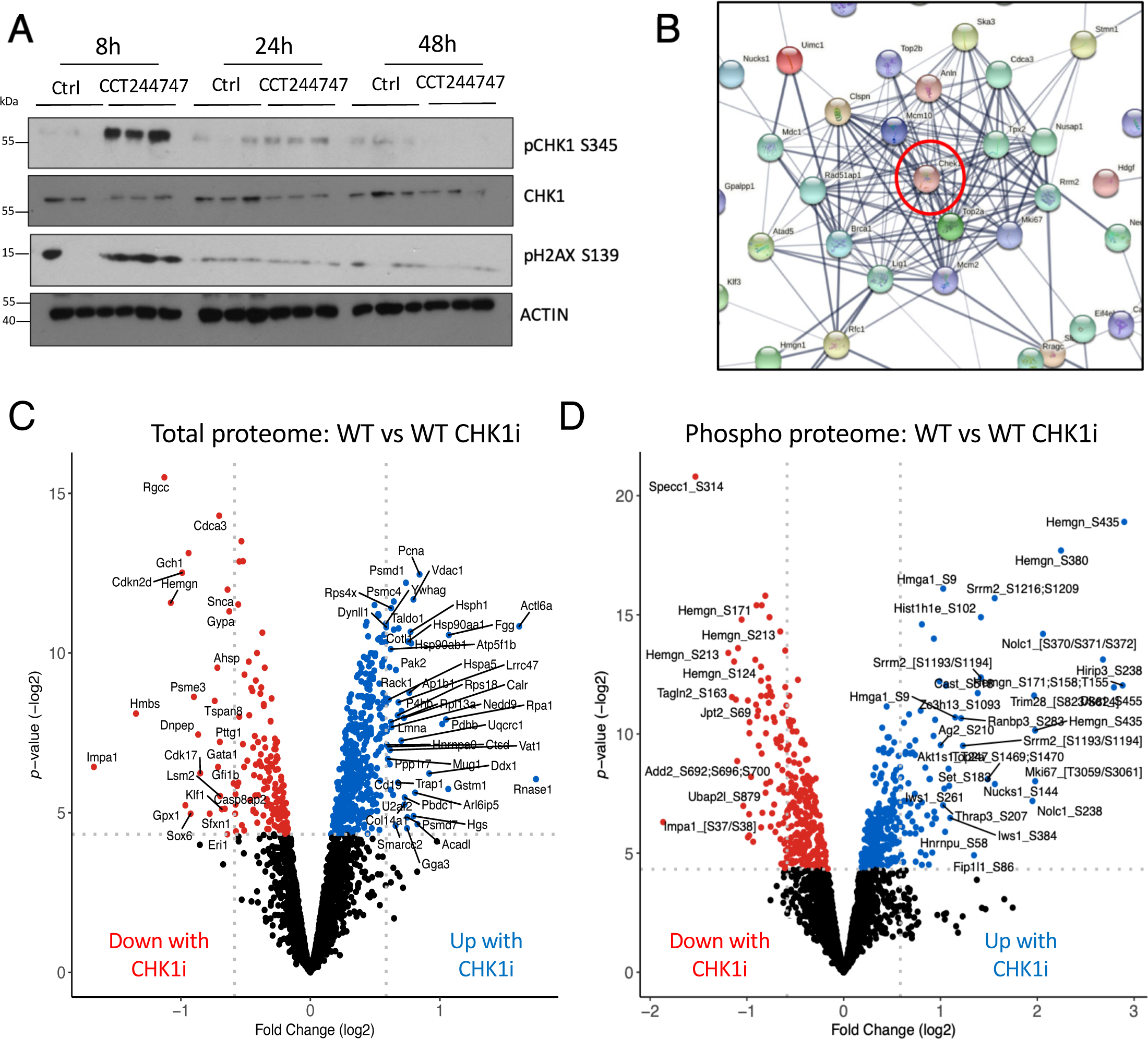
(A) Western blot analysis of phospho-Ser345 CHK1, CHK1, phospho-S139 H2AX (yH2AX) or ACTIN in snap frozen tumour extracts prepared from re-implanted Eµ-Myc and Eµ-Myc/*c-re/-/-* tumours mouse inguinal lymph nodes at given timepoints (8 hours, 24 hours or 48 hours) following a single dose of CCT244747. (B) STRING analysis showing that a cluster of proteins associated with CHK1 had down-regulated phosphorylation was upon CCT244747 treatment in Eµ-Myc WT tumours after a single dose of the CHK1i. (C & D) Volcano plots demonstrating a significant number of CCT244747 effects in Eµ-Myc WT lymphomas on both the total (C) and phospho (D) proteome, with both down- (red dots) and up-regulation (blue dots) being observed.

To explore regulation of phosphorylation-mediated signalling pathways in these re-implanted lymphomas, we used tandem mass tag (TMT)-based isobaric labelling to quantify relative changes in both total protein levels and phosphopeptide abundance (Fig S2B). Of the -4,000 proteins identified at a 1% false discovery rate (FDR), -2500 were quantified in at least three biological replicates (Supp Data File 1). At the phosphopeptide level, we identified over 6,500 phosphopeptides, quantifying -3,350 in at least three replicates (>4,500 in at least two bioreps; Supp Data File 1). STRING analysis (https://string-db.org/) of the phosphoproteomic data from WT Eµ-Myc lymphomas revealed a cluster of proteins known to be associated with CHK1, whose phosphorylation was down regulated upon CCT244747 treatment (Fig 2B, S2C, Supp Data File 2). This data confirmed effective targeting of CHK1 by CCT244747 *in vivo*.

Further analysis of this data demonstrated a significant number of CCT244747 effects in WT Eµ-Myc lymphomas, with 622 proteins and 625 phosphopeptides exhibiting a significant up- or down-regulation (P-value ::5 0.05) (Fig 2C & D). By contrast, relatively few effects of CHK1 inhibition were seen at the mRNA level from parallel RNA Seq data analysis (Fig S2D, Supp Data Files 3 & 4). This indicated that 8 hours after a single dose of CCT244747, the vast majority of the effects seen are post-transcriptional in nature. Strikingly, in comparison, relatively few significant changes were seen on the total and phospho proteomes following acute CCT244747 treatment of c-Rel^-/-^ Eµ-Myc lymphomas, with only 162 proteins and 89 phosphopeptides being significantly differentially regulated (P-value ::5 0.05) (Fig 3A & B, Fig. S3A). This was consistent with the lack of effectiveness on lymphoma growth seen with long term CCT244747 dosing (Fig. 1B & C).

**Figure 3.**
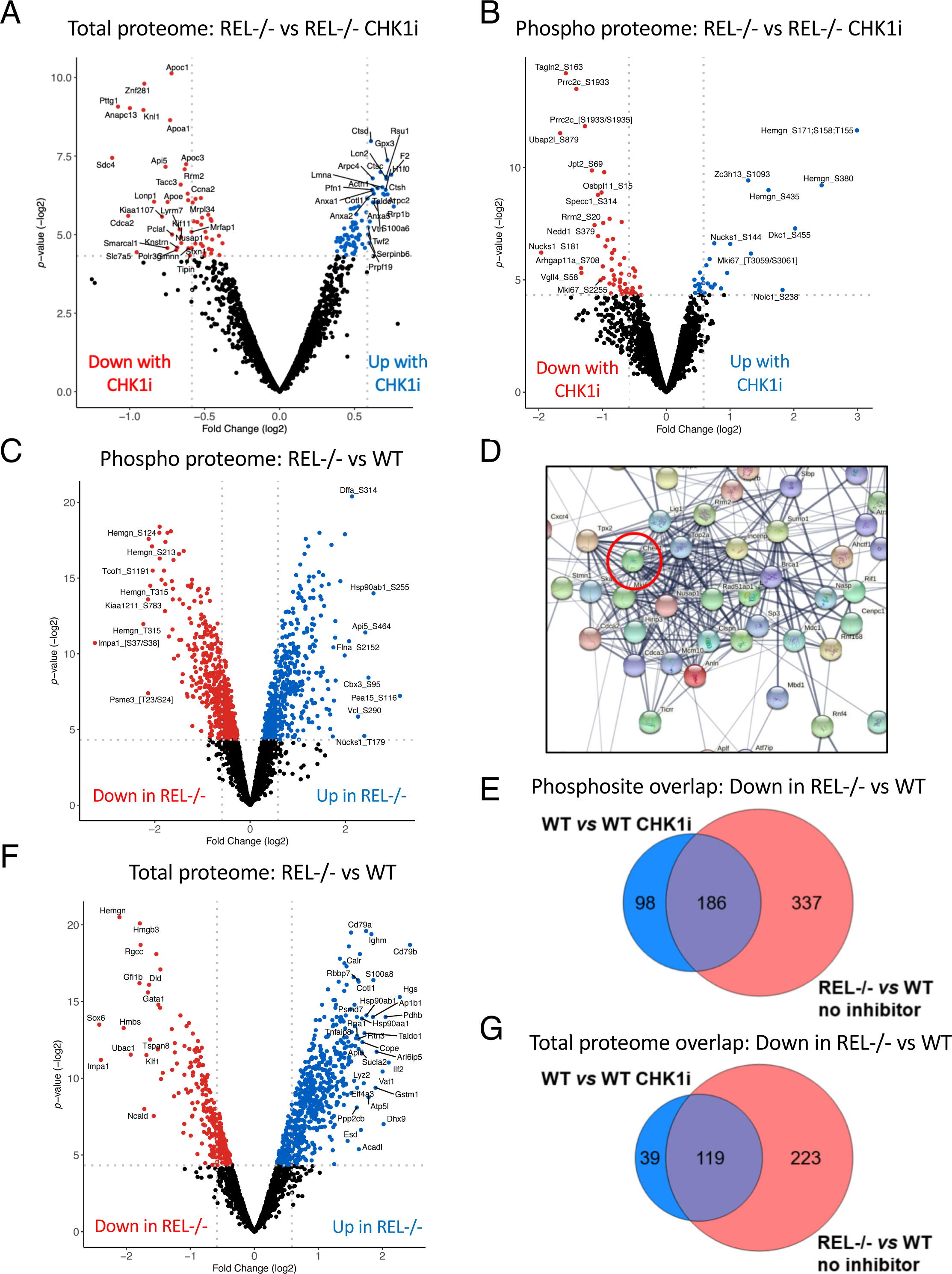
(A & B) Volcano plots illustrating that there was a much reduced number of CCT244747 effects in Eµ-Myc c-Rel^-/-^ lymphomas on both the total (C) and phospho (D) proteome. Down-regulation is shown with the red dots and up-regulation is shown with the blue dots. (C) Volcano plot demonstrating the significant number of phospho-proteomic differences between the Eµ-Myc WT and Eµ-Myc c-Rel^-/-^ lymphomas. 589 down-regulated phospho-peptides (shown by the red dots were observed) and 517 up-regulated phospho-peptides (blue dots) in Eµ-Myc c-Rel^-/-^ tumours when compared with Eµ-Myc WTs. (D) STRING analysis of the proteins associated with the 589 down-regulated phospho-peptides in the Eµ-Myc c-Rel^-/-^ lymphomas revealed that many of these had known linkages with CHK1 or CHK1 signalling. (E) Venn diagram illustrating that of the 284 unique downregulated phosphosites seen in Eµ-Myc WT tumours following acute CCT244747 treatment, 186 were also downregulated in c-Rel^-/-^ Eµ-Myc lymphoma cells without inhibitor treatment. (F) Volcano plot demonstrating the significant number of total protein differences between the Eµ-Myc WT and Eµ-Myc c-Rel^-/-^ lymphomas. Down-regulated proteins are shown with red dots and up-regulated proteins are shown with blue dots. (G) Venn diagram illustrating that of the 158 downregulated proteins seen in Eµ-Myc WT tumours following acute CCT244747 treatment, 119 were also downregulated in c-Rel^-/-^ Eµ-Myc lymphoma cells without inhibitor treatment.

### c-Rel -/- Eµ-Myc lymphomas have intrinsically downregulated the CHK1 pathway prior to inhibitor treatment

To better understand the underlying mechanistic basis that explains the relatively few significant (phospho)protein changes observed in c-Rel -/- Eµ-Myc lymphomas in response to treatment with CCT244747, we compared the protein and gene expression profiles of reimplanted WT and c-Rel^-/-^ Eµ-Myc lymphomas in the absence of CCT244747 treatment. The proteomic data revealed that c-Rel^-/-^ Eµ-Myc lymphoma cells had substantially rewired their cell signalling pathways, with a high level of both down and upregulated phosphopeptides compared to wild type (Fig. 3C, S3A). Furthermore, -75% of the protein level changes (and over 62% of the phosphorylation changes) that were induced in response to Chk1i in the WT Eµ-Myc lymphomas were also observed in the c-Rel^-/-^ samples with no treatment, suggesting that inhibition of Chk1 with CCT244747 may be working in part by modulating c-Rel-dependent processes (Supp Fig 3b &C, Supp Data File 5.

STRING analysis of proteins with downregulated phosphopeptides revealed that many have known connections to CHK1 (Fig 3D, Fig S3D, Supp Data File 2). Moreover, of the 284 unique downregulated phosphosites seen in wild type cells upon CCT244747 treatment, 186 (65%) were also downregulated in c-Rel^-/-^ Eµ-Myc lymphoma cells (Fig 3E, Supp Data File 5). Analysis of the total protein differences between reimplanted c-Rel^-/-^ Eµ-Myc lymphomas and their wild type counterparts, either with or without CCT244747 treatment revealed a similar trend. There were substantial differences between WT and c-Rel^-/-^ Eµ-Myc lymphomas in the absence of CHK1 inhibition (Fig 3F). Moreover, there was considerable overlap between these differences and the effect observed in WT cells upon treatment with CCT244747 also being downregulated in c-Rel^-/-^ Eµ-Myc lymphomas (without CHK1i treatment) (Fig 3G, Supp Data File 5). Interestingly, the magnitude of the changes seen in the CCT244747 treated wild type cells was generally lower in both cases than that seen constitutively in c-Rel^-/-^ Eµ-Myc lymphomas (Fig. S4). These results demonstrated that *c-Re/^-/-^* Eµ-Myc lymphomas have an intrinsic defect in CHK1 kinase signalling, comparable to the effect of inhibiting CHK1 in WT Eµ-Myc cells.

We next analysed our RNA Seq data to obtain further insights into the intrinsic transcriptional reprogramming of *c-Re/^-/-^* Eµ-Myc lymphoma cells that leads to these proteomic and cell signalling alterations (Supp Data Files 3 & 4). Functional profiling of the genes whose mRNA expression varied between WT and *c-Re/^-/-^* Eµ-Myc cells, revealed that of the 36 genes associated with ’Activation of ATR in response to replication stress’ (REAC:R-HSA-176187), 32 (89%) were down regulated in *c-Re/^-/-^* Eµ-Myc cells (Fig.S5A). This included transcript levels of CHEK1, which we subsequently validated using qPCR (Fig 4A). However, of the 342 proteins whose levels were decreased in *c-Re/^-/-^* Eµ-Myc cells compared to their wild-type counterparts, only 163 (47%) were also down regulated at the transcript level suggesting that there are very large post-transcriptional effects on protein expression (Fig S5B & Supp Data File 5).

**Figure 4.**
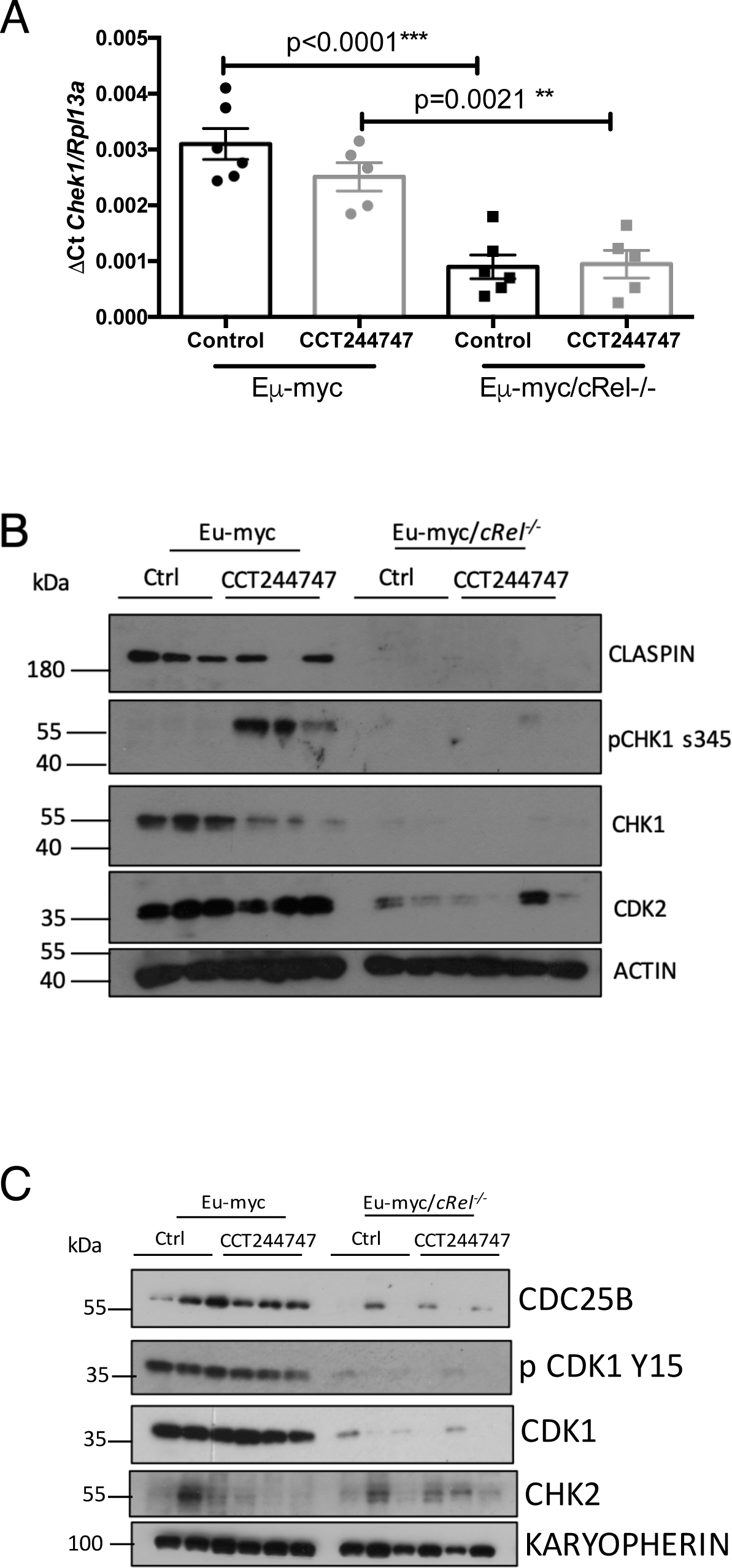
(A) Q-PCR validation of RNA-Seq analysis. Relative CHEK1 transcript levels are significantly reduced in tumours from Eµ-Myc/*c-re/-/-* (n=6) when compared with Eµ-Myc WTs (n=6). Data represents mean ± SEM. p** <0.01, p***<0.001 (One-way ANOVA with Tukey’s post-hoc test). CHEK1 expression is also partially reduced in WT tumours following CCT244747 treatment. Data represents mean ± SEM, each point is an individual mouse. (B) Western blot analysis of CLASPIN, phospho-Ser345 CHK1, CHK1, CDK2 or ACTIN in snap frozen tumour extracts prepared from re-implanted Eµ-Myc and Eµ-Myc/*c-re/-/-* tumours mouse inguinal lymph nodes 8 hours following a single dose of CCT244747. The expression of CHK1 and related pathway components are lost in Eµ-Myc/*c-re/-/-* tumours. (C) Western blot analysis of CDC25B, phospho-Tyr15 CDK1, CDK1, CHK2 or KARYOPHERIN in snap frozen tumour extracts prepared from re-implanted Eµ-Myc and Eµ-Myc/*c-re/-/-* tumours mouse inguinal lymph nodes 8 hours following a single dose of CCT244747. CDC25B, phospho-Tyr15 CDK1, CDK1, expression is lost in Eµ-Myc/*c-re/-/-* tumours, CHK2 is unaffected.

Western blot analysis confirmed not only that signalling through CHK1 was impaired in *c-Re/^-/-^* Eµ-Myc cells, but that there was almost complete loss of CHK1, CLSPN, CDC25B and CDK2 protein (Figs 4B & C, and S5C). Taken together, these data suggest that the *de novo* resistance of the *c-Re/^-/-^* Eµ-Myc lymphoma cells to CCT244747 arises from these cells already having downregulated the CHK1 pathway. Consequently, further attempts to inhibit CHK1 have little effect.

### *A*cquired resistance to CHK1 inhibition in U20S cells is also associated with downregulation of CHK1 protein

We next wished to determine if similar effects on CHK1 levels and activity were seen as a consequence of acquired CHK1 inhibitor resistance. To this end, we generated four independent isolates of the osteosarcoma cell line, U20S, with resistance to the CHK1i, CCT244747. This was achieved through long term culture in increasing concentrations of CCT244747. Eventually, the resistant U20S cells were able to proliferate in high CCT244747 concentrations (Fig 5A), and retain clonogenic potential (Fig 5B). As controls we also passaged U20S cells in the absence of CCT244747 to mimic effects of long-term culture.

**Figure 5.**
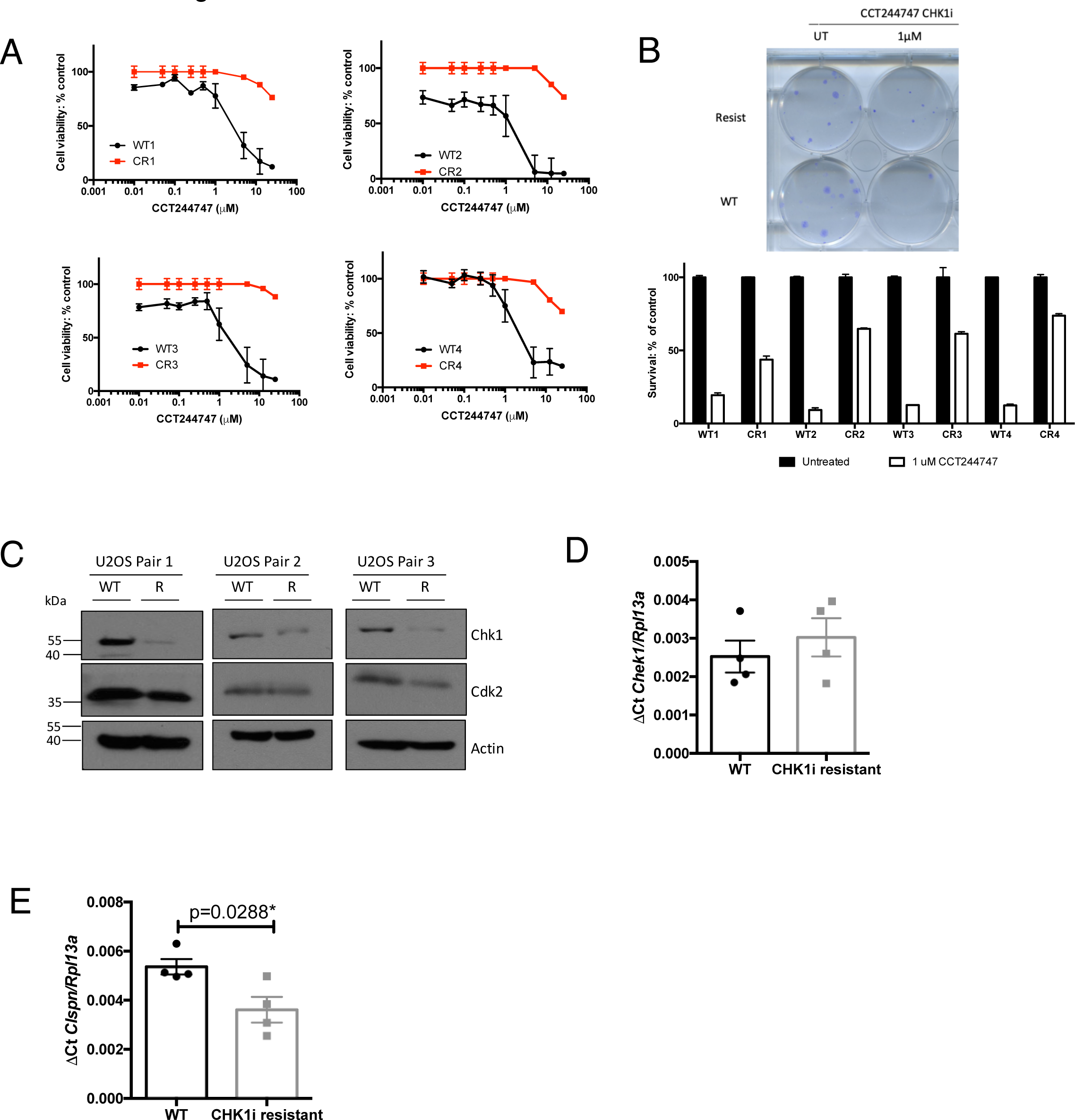
(A) Four independently derived CCT244747 resistant (CR) U20S cell lines are resistant to CHK1 inhibitor treatment. Cell viability (Prestoblue assay) in WT and CR U20S following treatment with increasing concentrations of the CHK1 inhibitor, CCT244747 for 72 hours. (B) Increased clonogenic survival in four independently derived CCT244747 resistant (CR) U20S cell lines following CHK1 inhibitor treatment. Representative image and bar graph data showing clonogenic survival in WT and CR U20S following either treatment with 1 μM CCT244747for 24 hours. (C) Western blot analysis of CHK1, CDK2, or ACTIN in extracts prepared from WT and CCT244747 resistant (CR) U20S. (D) Q-PCR data showing relative Chek1 expression in four independently derived CCT244747 resistant (CR) U20S cell lines, or their WT counterparts. Chek1 expression is unaffected in CR U20S. Data represents mean ± SEM, each point is the mean of three independent experiments in each of the four cell lines. (E) Q-PCR data showing relative Claspin expression in four independently derived CCT244747 resistant (CR) U20S cell lines, or their WT counterparts. Claspin expression is reduced in CR U20S. Data represents mean ± SEM, each point is the mean of three independent experiments in each of the four cell lines p* <0.05 (Unpaired Student’s t-test).

To determine whether CHK1 signalling was affected during the acquisition of resistance, we performed Western blotting and qPCR analyses. Western blot analysis confirmed that CHK1 levels were reduced in three out of four CCT244747 resistant isolates (Fig 5C & 6E). We also observed a reduction in Claspin transcript levels in CHK1i resistant U20S cells (Fig 5D), mirroring the observations in our resistant mice (Fig 1A). However, by contrast with our data from *c-Re/^-/-^* Eµ-Myc lymphoma cells, there was no reduction in CHK1 mRNA levels as determined by RNA Seq and qPCR analysis (Fig. 5D, Supp Data Files 6 & 7). We also failed to observe a reduction in expression of the 32 genes associated with ’Activation of ATR in response to replication stress’ (> 2 fold change, adj p-value <0.05; REAC:R-HSA-176187) that were down regulated in the *c-Re/^-/-^* Eµ-Myc cells. The exception to this was again CLSPN, where the RNA Seq data confirmed a 2.2 fold downregulation (adj p-value = 0.0037) in CCT244747 resistant U20S cells (Fig 5E & Supp Data Files 6 & 7).

Taken together, the data suggest a clear and consistent mechanism of both *de novo* and acquired resistance, namely down regulation CHK1 levels and thus activity, thereby rendering cells insensitive to a CHK1 inhibitor.

### Deregulation of ubiquitin mediated proteolysis in *c-Rel^-/-^* Eµ-Myc lymphoma cells

Results from our Eµ-Myc lymphoma cell proteomic analysis above, together with the loss of CHK1 protein but not mRNA in the CCT244747 resistant U20S cells, suggested that post-transcriptional regulation of protein levels was also a key factor in acquisition of both *de novo* and acquired CHK1 inhibitor resistance. We therefore further analysed our RNA Seq data from wild type and *c-Re/^-/-^* Eµ-Myc lymphoma cells. 0f the genes associated with ubiquitin dependent proteolysis we observed a number of changes. Most strikingly, there was significant down regulation of the deubiquitinases (DUBs) USP1 and USP14 (Fig. 6A, Supp Data Files 3 & 4). USP1 has been reported as a key regulator of DNA repair and is known to play a role in stabilising members of the DNA damage response, such as FANCD2 and PCNA [27, 28] by removing the K48 ubiquitin degradation signal. Interestingly, one report suggested that USP1 can act directly as a DUB for CHK1, by protecting it from proteasomal degradation [29], USP14 is often overexpressed in tumours and has been shown to deubiquitinate and stabilise the androgen receptor in models of breast and prostate cancer [30, 31]. Downregulation of these genes was validated by qPCR (Fig 6B), while western blot analysis revealed almost total loss of these proteins in extracts prepared from *c-Rel^-/-^* Eµ-Myc lymphoma cells (Fig. 6C & S6A). Moreover, both USP1 mRNA and protein were down regulated in U20S CCT244747 resistant cells (Fig. 6D & E). siRNA depletion of c-Rel in wild type U20S cells, resulted in a reduction of both USP1 and USP14 mRNA and protein and this was associated with downregulation of CHK1 protein but not CHK1 mRNA (Fig. 6F). This suggests a conserved mechanism through which c-Rel can directly or indirectly control the transcription of USP1 and USP14, with the loss of one or both of these DUBs then results in CHK1 protein destabilisation.

**Figure 6.**
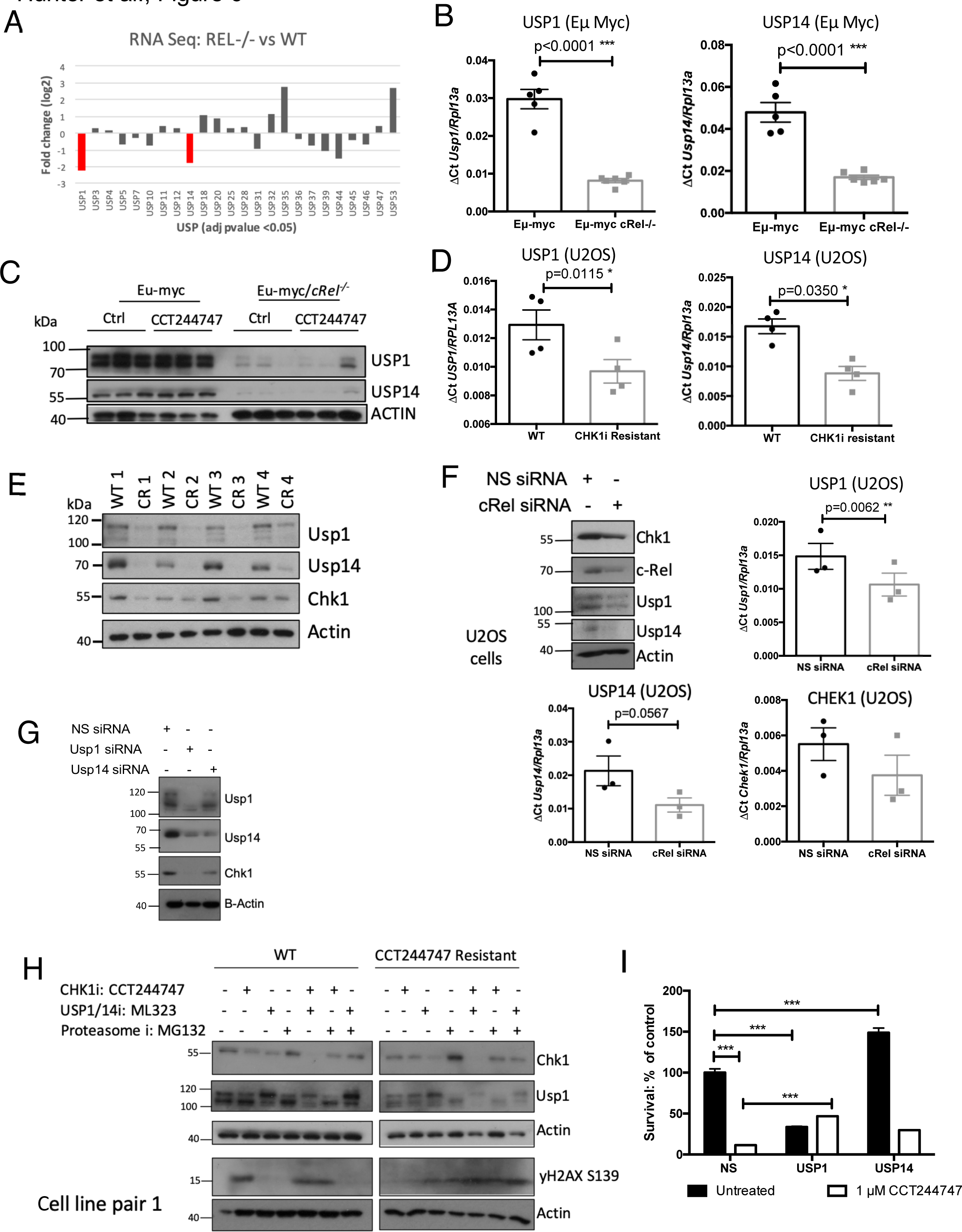
(A) Bar graph showing the relative expression of 24 DUBs that were significantly up- or down-regulated in the Eµ-Myc/*c-re/-/-* tumours by RNA-Seq analysis. The red bars show that both USP1 and USP14 were down-regulated by approximately 2 fold compared with Eµ-Myc WTs. (B) Q-PCR validation of RNA-Seq analysis. Relative USP1 and USP14 transcript levels are significantly reduced in tumours from Eµ-Myc/*c-re/-/-* (n=6) when compared with Eµ-Myc WTs (n=5). Data represents mean ± SEM. p** <0.01, p***<0.001 (Unpaired student’s t-test). Data represents mean ± SEM, each point is an individual mouse. (C) Western blot analysis of USP1, USP14 or ACTIN in snap frozen tumour extracts prepared from re-implanted Eµ-Myc and Eµ-Myc/*c-re/-/-* tumours mouse inguinal lymph nodes 8 hours following a single dose of CCT244747. USP1 and USP14 expression is lost in Eµ-Myc/*c-re/-/-* tumours. (D) Q-PCR data showing relative USP1 and USP14 transcript levels are significantly reduced in in four independently derived CCT244747 resistant U20S cell lines, compared with WT U20S cells. Data represents mean ± SEM. p* <0.05 (Unpaired student’s t-test). Data represents mean ± SEM, each point is the mean of three independent experiments in each of the four cell lines. (E) Western blot analysis of USP1, USP14, CHK1, or ACTIN in extracts prepared from WT and CCT244747 resistant U20S. USP1 and USP14 expression is lost in CCT244747 resistant U20S. (F) Western blot and Q-PCR analysis from WT U20S cells following siRNA targeting c-Rel or a Non-specific siRNA control. Western blot analysis shows that knockdown of c-Rel results in a reduction of USP1, USP14 and CHK1. ACTIN is used as a loading control. Q-PCR data shows that USP1 and USP14 transcript levels are reduced following c-Rel knockdown, but that CHEK1 transcript levels are unaffected. Data represents mean ± SEM, each point is the mean of three independent experiments. p** <0.01 (Unpaired student’s t-test). (G) Western blot analysis from WT U20S cells following siRNA targeting USP1, USP14 or a Non-specific siRNA control. Data shows that CHK1 is completely lost following USP1 knockdown and partially lost following USP14 knockdown. ACTIN is used as a loading control. (H) Western blot analysis of WT or CCT244747 resistant U20S cells treated with CCT244747, the USP1/14 inhibitor, ML323, or the Proteasome inhibitor MG-132, alone or in combination. Blots were probed for CHK1, USP1, yH2AX or ACTIN. Inhibition of USP1/14 in WT U20S results in the loss of CHK1. Proteasomal inhibition in the CCT244747 resistant U20S cells results in the stabilization of CHK1 protein. (I) Clonogenic survival in WT U20S cell lines following siRNA targeting USP1, USP14 or a Non-specific siRNA control. U20S are sensitive to CCT244747 in the presence of control or USP14 siRNA, however knockdown of USP1 renders them insensitive to CCT244747 treatment. Data represents mean ± SEM, each point is the mean of three independent experiments. p*** <0.01 (0ne-way AN0VA with Tukey’s post-hoc test).

To determine whether USP1, or USP14 were responsible for CHK1 protein stability in our model, we depleted both with siRNAs in WT U20S cells. We found that loss of USP1 resulted in almost total loss of CHK1 at the protein level, suggesting that in the absence of this DUB, CHK1 is targeted by the proteasome for degradation (Fig. 6G). Depletion of USP1 also reduced USP14 levels, while the USP14 siRNA resulted in partial loss of CHK1, suggesting that the activity of these DUBs may be linked. This was confirmed by treatment of WT U20S cells with the USP1 inhibitor ML323 [32], which also resulted in a reduction in CHK1 protein levels (Fig. 6H and S6B). 1mportantly, inhibition of the proteasome using MG132, restored CHK1 protein in the resistant U20S cells (Fig 6H and S6B), and this in turn induced a DNA damage response indicated by an increase in yH2AX phosphorylation, suggesting a potential restoration of CHK1i sensitivity.

To determine whether loss of either USP1 or USP14, was responsible for the resistance to CHK1 inhibitors we performed clonogenic assays following knockdown of either USP1 or USP14 in combination with CCT244747 treatment. Although, loss of USP1 itself reduced the clonogenic potential of U20S cells, the remaining cells now exhibited complete resistance to CHK1 inhibition (Fig. 6I). By contrast, depletion of USP14 U20S cells had little effect on either clonogenic potential or CCT244747 sensitivity (Fig 6I). These data indicates that loss of USP1 can contribute to CHK1 inhibitor resistance in this model.

### Pl3K/AKT signalling is upregulated in CHK1i resistant lymphoma cells, and provides an alternative therapeutic strategy

The dramatic loss of CHK1 signalling we observed in the *c-Re/^-/-^* Eµ-Myc lymphoma cells raised the question of how these cells were surviving and coping with ongoing DNA replication stress. Especially given our previous observation [24] that lymphomas in these mice arise earlier than in wild type controls. The activation of compensatory signalling pathways is a major contributing factor in the development of resistance to kinase inhibitors [33–38], and often goes hand-in-hand with the tumour cell having bypassed the need for the drug target, which we observed in our models of both *de novo* and acquired resistance (Figs 4 & 5).

USP1 has also been demonstrated to be a negative regulator of AKT kinase signalling [39] We therefore further analysed our proteomic analysis following acute CHK1i treatment (Fig 2) to see if there was any evidence for altered AKT pathway activity, focusing this time on upregulated phosphosites and proteins. These analyses revealed 480 upregulated phosphosites (p<0.05) and 624 upregulated proteins (p<0.05) in c-Rel^-/-^ Eµ-Myc lymphomas with no CCT244747 treatment compared to wild type controls (Fig. 7A & B, S3A, Supp Data File 5). Similar to our observation with downregulated phosphosites and proteins, there was considerable overlap with those seen in wild type Eµ-Myc lymphomas after CCT244747 treatment. Of the 294 phosphosites and 464 proteins elevated in wild type Eµ-Myc lymphomas in response to CCT244747, 51% and 74% respectively were also upregulated in c-Rel^-/-^ Eµ-Myc cells (Fig. 7A & B, Supp Data File 5). However, the magnitude of the increases seen in the CCT244747 treated wild type cells was lower in both cases than that seen constitutively in c-Rel^-/-^ Eµ-Myc lymphomas (Fig. S4). Nonetheless, this further confirmed the similarities between c-Rel^-/-^ Eµ-Myc lymphomas and their wild type equivalents following CHK1 inhibition.

**Figure 7.**
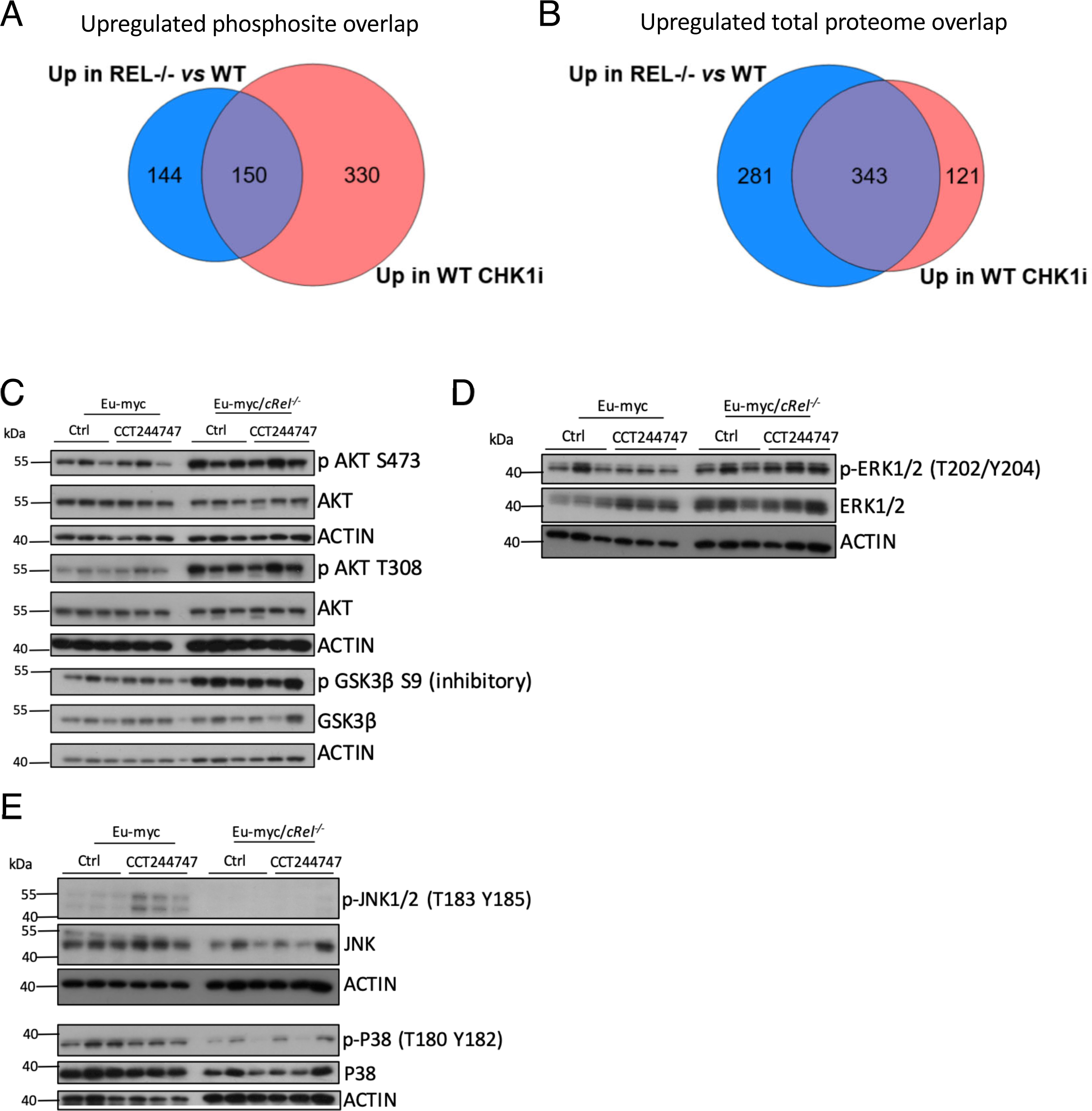
(A) Venn diagram illustrating that of the 294 unique upregulated phosphosites seen in Eµ-Myc WT tumours following acute CCT244747 treatment, 150 were also upregulated in c-Rel^-/-^ Eµ-Myc lymphoma cells without inhibitor treatment. (B) Venn diagram illustrating that of the 464 upregulated proteins seen in Eµ-Myc WT tumours following acute CCT244747 treatment, 343 were also upregulated in c-Rel^-/-^ Eµ-Myc lymphoma cells without inhibitor treatment. (C) Western blot analysis of phospho-Ser473 AKT, phospho-Thr308 AKT, AKT, phospho-Ser9 GSK3B, GSK3B or ACTIN in snap frozen tumour extracts prepared from re-implanted Eµ-Myc and Eµ-Myc/*c-re/-/-* tumours mouse inguinal lymph nodes 8 hours following a single dose of CCT244747. The data shows that the AKT pathway is highly active in Eµ-Myc/*c-re/-/-* tumours. (D) Western blot analysis of phospho-Thr202/Tyr204 ERK1/2, ERK1/2 or ACTIN in snap frozen tumour extracts prepared from re-implanted Eµ-Myc and Eµ-Myc/*c-re/-/-* tumours mouse inguinal lymph nodes 8 hours following a single dose of CCT244747. The data shows that the ERK pathway is highly active in Eµ-Myc/*c-re/-/-* tumours. (E) Western blot analysis of phospho-Thr183/Tyr185 JNK1/2, JNK1/2, phospho-Thr180/Tyr182 p38, p38 or ACTIN in snap frozen tumour extracts prepared from re-implanted Eµ-Myc and Eµ-Myc/*c-re/-/-* tumours mouse inguinal lymph nodes 8 hours following a single dose of CCT244747. The data shows that the JNK/p38 pathway signalling is reduced in Eµ-Myc/*c-re/-/-* tumours.

Given the links between USP1 and AKT, we next examined potential links with known AKT networks. STRING analysis revealed that 71 phosphorylated proteins upregulated in c-Rel^-/-^ Eµ-Myc lymphomas had known links to AKT1 (Fig. S7, Table S1, Supp Data File 2). This included AKT1S1, and analysis of the phosphoproteomic data (Supp Data File 1) revealed this was a consequence of phosphorylation at T247, a known site of AKT (Protein Kinase B) phosphorylation [40]; To determine if these upregulated phosphoproteins arose due to changes in AKT activity, we examined AKT1 phosphorylation status by western blot, revealing significant upregulation of AKT1 phosphorylation on Ser473 and Thr308 in the c-Rel^-/-^ lymphomas, phosphorylation sites know to be required for full AKT activity (Fig 7C, S8. lnterestingly, the AKT1 activator, Pl3K, also exhibited elevated levels of phosphorylation P85 (Y458) / P55 (Y199) (Fig S8). AKT phosphorylation is known to induce a pro-survival response by inducing the phosphorylation of GSK3β on Ser9, which inhibits its activity, and this again was upregulated in the c-Rel^-/-^ cells. We also observed an increase in ERK 1/2 phosphorylation at its activation sites T202/Y204. but not of JNK or p38 MAP kinase signalling (Fig 7C-E, S8). This result potentially explains how these tumours are surviving loss of CHK1, as they have hyper-activation of the Pl3K/AKT and ERK pathways, which are known to inhibit apoptosis and cell cycle arrest [41, 42].

We hypothesised that if upregulation of the Pl3K/AKT and ERK pathways allowed c-Rel^-/-^ Eµ-Myc lymphomas to survive in the absence of CHK1, then targeting either of these pathways would be a good therapeutic strategy. We decided to evaluate the effectiveness of the Pl3K inhibitor, GDC-0941/Pictilisib [43] *in vivo,* by analysing its effect on the growth of three transplanted WT Eµ-Myc and Eµ-Myc/*c-Re/^-/-^* tumours. As previously performed with CCT244747 (Fig. S1A), each tumour was implanted into six syngeneic C57Bl/6 recipient mice and three were treated orally with GDC-0941 once a day for nine days, while three received a vehicle control (Fig. 8, S9 & 10). After treatment, we observed a striking reduction in lymphoid tumour burden in all mice re-implanted with c-Rel^-/-^ Eµ-Myc tumours and treated with GDC-0941 (Figs Fig. 8, S9 & 10). By contrast, in all of the WT Eµ-Myc tumours, no significant reduction in lymphoid tumour burden was seen after GDC-0941 treatment. These data confirmed that elevated Pl3K/AKT signalling critically compensates for loss of c-Rel^-/-^ in these Eµ-Myc lymphomas.

**Figure 8.**
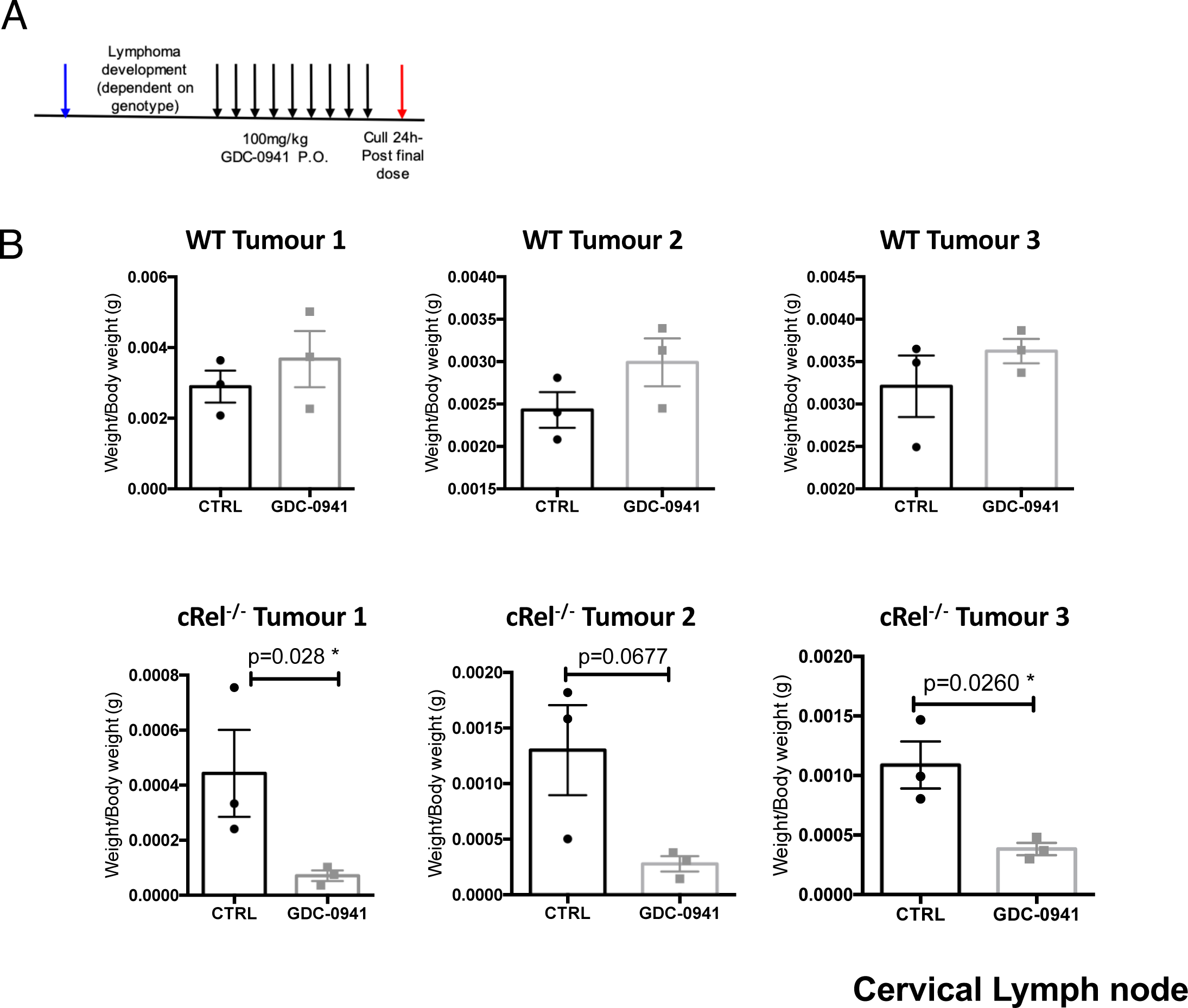
(A) Schematic diagram illustrating the PI3Ki *in vivo* study in Eµ-Myc and Eµ-Myc/*c-re/-/-* mice. 6 week old C57Bl/6 WT mice were implanted with either Eµ-Myc or Eµ-Myc/*c-re/-/-* (blue arrow) and once tumours became palpable were treated with either 100 mg/kg GDC-0941/Pictilisib p.o or vehicle control once daily for 9 days (black arrows). Mice were euthanised 24 hours after the final dose (red arrow) and tumour burden assessed. (B) Scatter showing the response of three reimplanted Eµ-Myc (top panel) and three Eµ-Myc cRel^-/-^ (bottom panel) tumours and their response to GDC-0941/Pictilisib in the cervical lymph node tumour site. Each of the 3 tumours was implanted into 6 syngeneic recipient C57Bl/6 mice, 3 were treated with GDC-0941/Pictilisib (100 mg/kg p.o), and 3 with vehicle control, for 9 days once tumours became palpable. A response was defined as a significant reduction (or increase) in tumour burden (P<0.05) using unpaired Student’s t-tests. WT Eµ-Myc showed little response to GDC-0941/Pictilisib whereas the Eµ-Myc cRel^-/-^ tumours were reduced by GDC-0941/Pictilisib.

## Discussion

Loss of c-Rel has many effects in the Eµ-Myc lymphoma model, underlining the critical role this NF-κB subunit plays in this context [24]. Indeed, our proteomic data demonstrates that these *c-Re/^-/-^* Eµ-Myc lymphoma cells undergo a comprehensive rewiring of their cell signalling pathways. Here we have explored the basis for the resistance of *c-Re/^-/-^* Eµ-Myc lymphomas to CHK1 inhibition, and revealed a pathway regulating the response to DNA replication stress in cancer.

One dramatic finding underpinning the development of resistance to the CCT244747 CHK1 inhibitor in this model, is the loss of CHK1 protein itself, together with other components of the DNA replication stress response and cell cycle pathways. The mechanisms underlying this effect are complex but at the core of this is down regulation of the mRNA and protein of the USP1 DUB in *c-Re/^-/-^* cells. USP1 has been previously linked to numerous components of the DNA damage response [27, 28, 44] and implicated as a CHK1 DUB [29]. We propose that c-Rel directly, or indirectly, regulates the expression of USP1. Consequently, in the absence of c-Rel, loss of USP1 results in destabilisation of CHK1 and other pathway components (Figure 9). Contributing towards this effect, is the parallel loss of Claspin expression (Figure 1D, 4B, 5E). Whether it is the initial loss of Claspin, thus breaking the link between ATR and CHK1, that leads to subsequent effects via USP1 is not known. Nonetheless, it is probable that the parallel loss of USP1 and Claspin works synergistically to down regulate CHK1 protein levels and activity. This complexity, however, ultimately creates a simple explanation for CHK1 inhibitor resistance in the *c-Re/^-/-^* Eµ-Myc lymphoma cells: the target of the drug is no longer present.

**Figure 9.**
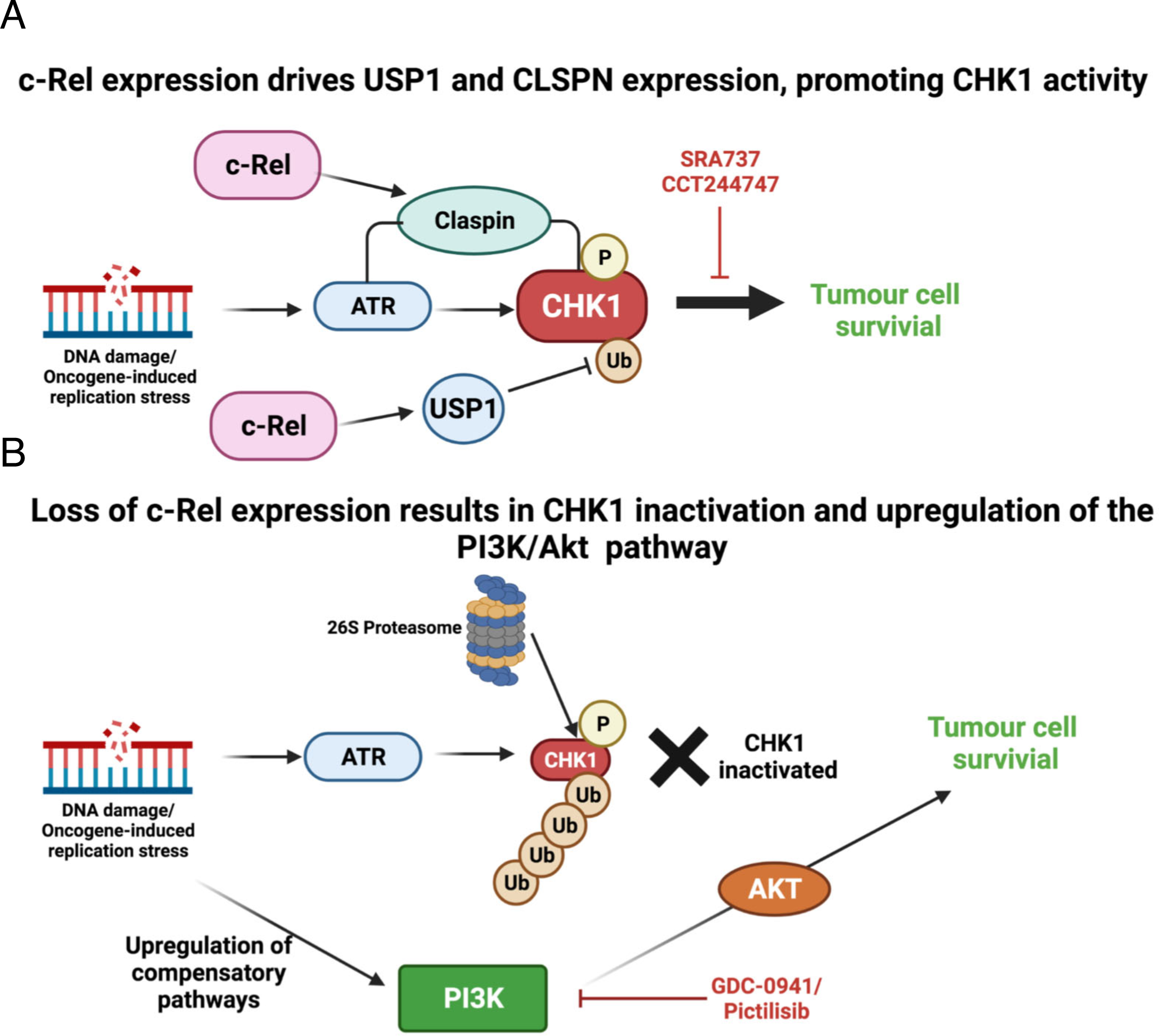
MYC-driven tumours, or those undergoing high levels of replication stress become addicted to ATR-CHK1 signalling. When cRel in present and controlling the levels of the adaptor protein Claspin, which facilitates the phosphorylation of CHK1 by ATR, and also the levels of USP1, the known CHK1 DUB, this addiction to CHK1 signalling renders the cell sensitive to CHK1 inhibition by CCT244747 or SRA737, leading to tumour cell death. The data herein indicates that both *de novo* and acquired resistance to CHK1i is a two-step process. First there is loss of CHK1 protein, or pathway activity, this can be due to the loss of Claspin or USP1, thus rendering the cell insensitive to CHK1 inhibition. Secondly, we observe the activation of compensatory pathways such as the Pl3K/AKT and ERK pathways to allow the tumour cell to survive the loss of CHK1. This can be targeted with agents such as the Pl3K inhibitor, GDC-0941/Pictilisib to result in tumour cell death.

Our proteomic data demonstrates that *c-Re/^-/-^* Eµ-Myc lymphomas strikingly resemble wild type Eµ-Myc cells that had received the CHK1 inhibitor (Figure 3). This raised the question of why did the WT cells die when CHK1 is inhibited but the *c-Re/^-/-^* Eµ-Myc lymphoma do not. We observed significant upregulation of phospho and total proteins in both WT Eµ-Myc lymphomas treated with the CHK1 inhibitor as well as intrinsically in *c-Re/^-/-^* cells (Figures 2, 3 & 7). This suggested that activation of a compensatory pathway was allowing the *c-Re/^-/-^* Eµ-Myc lymphomas to survive and proliferate, even in the absence of CHK1 and the presence of ongoing DNA replication stress. Since USP1 has also been demonstrated to be a negative regulator of AKT pathway signalling [39, 45–47], we took a candidate approach to resolving this. Examination of the phospho proteomic data and subsequent western blot analysis of protein extracts, revealed that indeed, AKT1 pathway activity was upregulated in *c-Re/^-/-^* Eµ-Myc lymphomas (Figure 7). Moreover, treatment with the Pl3K inhibitor Pictilisib, selectively targeted the *c-Re/^-/-^* Eµ-Myc lymphomas, while having no effect on their WT equivalents (Figure 8). The second component of our model therefore is that loss of c-Rel and consequently USP1, automatically results in activation of a parallel survival pathway involving Pl3K and AKT. lmportantly, in the phospho proteomic data, we also observed similar upregulation of AKT1 linked phospho proteins in WT Eµ-Myc lymphomas treated with CCT244747, albeit at a much reduced level relative to the *c-Re/^-/-^* Eµ-Myc lymphomas (Supp Figure 7). This implies that cotargeting of Pl3K or AKT might increase the efficiency of CHK1 inhibition in cancer therapy.

This study does not address how relevant this data is to human cancer in general and the use of CHK1 inhibitors clinically. However, key components of the work from Eµ-Myc lymphomas, such as down regulation of CHK1 and USP1 protein, were also seen in independently derived populations of CCT244747 resistance U20S cells (Figure 6). Moreover, down regulation of USP1 in WT U20S cells resulted in loss of CHK1 protein and CCT244747 resistance. Nonetheless, the more widespread significance of these findings requires further study before it can be inferred that this is a common mechanism in human cancer.

Previous CHK1 inhibitor studies have shown other potential routes of drug sensitivity and resistance. An analysis of MK-8776, performed in multiple sensitive cell lines and resistant cell lines showed that cells responsive to this CHK1 inhibitor upregulated CDK2 and cyclin A after treatment, with an associated increase in double stranded DNA breaks [48]. Drug sensitivity was also associated with an increased accumulation of CDC25A [48]. By contrast, cells that were MK-8776 resistant failed to dephosphorylate and thus activate CDK2 [48]. Another study into the MK-8776 CHK1 inhibitor found that cells deficient in MRE11 were resistant to CHK1 inhibition [49]. A study of the CHK1 inhibitor LY2603618 (Rabusertib) in head and neck cell lines also found that sensitivity to the drug was dependent on CDK activity, and highlighted CDK1 as being significant in drug response. Here, increased CDK1 levels were indicative of reduced drug sensitivity, potentially due to upregulation of origin firing and this overcoming S phase replication stalling [50]. Research into the PF-00477736 CHK1 inhibitor resistant mantle cell lymphoma cell line JEK0-1, showed that resistant cells had a shorter S phase and a reduced expression of cell cycle checkpoint proteins, including cyclin D1 [51]. These studies both contrast with our analysis, where we found that *c-Re/^-/-^* Eµ-Myc lymphomas exhibited reduced levels of CDK1 and CDK2.

Given these contrasting findings, it is likely that the mechanism used in the development of CHK1 inhibitor resistance will be dependent on the tumour context, with both the cell type and oncogene/tumour suppressor status having a key role. Nonetheless, our data suggests that down regulation or mutation of USP1 is likely to be a common feature arising in patients undergoing therapy involving a CHK1 inhibitor. Not only will loss of USP1 lead to loss of the target of the inhibitor but will also result in upregulation of AKT or ERK activity as a compensatory mechanism. We propose that should CHK1 inhibitors become commonly used cancer therapeutics that this pathway of resistance be investigated further. 0ur data implies that treatment with a Pl3K or AKT pathway inhibitor could be a valuable strategy either in the treatment of patients whose tumours develop resistance to CHK1 inhibition or potentially as a combination therapy to enhance the effectiveness of initial treatment.

## Acknowledgements

We would like to thank lglika lvanova, Sonia Rocha, Laura Greaves, Niall Kenneth, Urszula McClurg, Suzanne Madgwick, Adrian Yemm, and all members of the NDP laboratory for helpful advice and assistance. JEH and AEC are funded by Cancer Research UK grant C1443/A22095. JEH previously received funding from Leukemia Lymphoma Research grant 11022 and Cancer Research UK grant C1443/A12750. NLH was funded by Cancer Research UK Clinical PhD studentship. JAB and HS were funded by Wellcome Trust grant 094409. lC and MDG receive funding from Cancer Research UK grant number C309/A11566, and MDG also from the University of Kent. This work was also supported by instrumentation funding from the Biotechnology and Biosciences Research Council (BBSRC; BB/M012557/1 and BB/R000182/1). Figure 9 was created with Biorender.com.

## Author contributions

JEH: performed majority of experimental work. Contributed to design of experiments and manuscript writing.

AEC: performed all proteomics work and analyses. Contributed to manuscript writing.

PJB: supported mass spectrometry methods development and proteomics data analysis.

NLH & SK: performed downstream analysis of mouse tissues.

SL, JAB & HS assisted with procedures involving Eµ-Myc.

RM: contributed to *in vitro* experimental work.

HDT: provided training and assisted with lymphoma re-implantation studies.

JC & LT: performed RNA-Seq analysis

PL & MSRH: performed bioinformatics analysis of RNA-Seq data

MDG and lC: contributed to the design and analysis of experiments.

CEE: contributed to design of experiments and manuscript writing.

NDP: contributed to design of experiments, analysis of RNA Seq and proteomic data, manuscript writing.

## Methods

### Ethics statement

All mouse experiments were approved by Newcastle University’s Animal Welfare and Ethical Review Board. All procedures, including the of breeding genetically modified mice, were carried out under project and personal licenses approved by the Secretary of State for the Home Office, under the United Kingdom’s 1986 Animal (Scientific Procedures). Animals were bred in the Comparative Biology Centre, Newcastle University animal unit, according to the FELASA Guidelines.

### Mouse models

*c-Rel^-/-^* mice were provided by Dr Fiona Oakley (Newcastle University) and Eµ-Myc mice were purchased from The Jackson Laboratory, Maine, USA. C57Bl/6 mice used for re-implantation studies were purchased from Charles River (UK). Male Eµ-Myc transgenic mice that were used as breeding stock were omitted from the survival analysis. In all experiments, the relevant pure C57Bl/6 (WT) strain was used as a control. No blinding of groups in mouse studies was performed. All mice were designated to an experimental group dependent on their genotype.

### Drugs and compounds

CCT244747 was synthesized as described [52] by MedKoo Biosciences. GDC-0941/Pictilisib was purchased from Med Chem Express. All other compounds were purchased from Sigma Aldrich.

### Resistant cell line generation

U2OS cells were cultured in increasing concentrations of CCT244747, starting with the IC_50_ concentration of 1 µM. The concentration of CCT244747 was doubled at each passage to a final concentration of 8 µM. WT controls were given an equivalent amount of DMSO to account for any DMSO-related toxicity.

### siRNA Knockdown transfections

U20S cells were transfected with 10 nM siRNA targeting either USP1, USP14 or cRel (0N-TARGET plus Smart pool, Dharmacon) or a Non-specific siRNA control using Dharmafect 4 transfection reagent, according to manufacturer’s protocols. Cells were harvested, or used in downstream assays 72 hrs post-transfection, once target depletion had been confirmed.

### Cell viability assays

Freshly isolated Eµ-Myc or Eµ-Myc/*c-Re/*^-/-^ tumour cells (5x10^5^ per well), or WT or CHK1i resistant U20S (5x10^3^ per well) were seeded into 96-well plates. Increasing concentrations of CHK1 inhibitor, CCT244747, or solvent controls were added to three replicate wells. After 96hrs, viability was quantified using the PrestoBlue Cell Viability Reagent (ThermoFisher Scientific, UK), according to manufacturer’s instructions.

### Cell survival assays

Exponentially growing WT or CHK1i resistant U20S were treated for 24h with 1 μM CHK1 inhibitor, CCT244747 or solvent controls before re-seeding onto Petri dishes at known cell number (1000, 2500 or 5000 cells/dish). Colonies were fixed 14 days later with methanol:acetic acid (3:1) and stained with 0.4% (w/v) Crystal Violet. Cloning efficiencies were normalized to untreated controls.

### Gene expression analysis using quantitative real-time PCR

Total RNA was purified from snap frozen Eµ-Myc or Eµ-Myc/*c-Re/*^-/-^ tumour samples by homogenisation using Precellys 24 ceramic mix bead tubes (Stretton Scientific Ltd) in a Precellys 24 benchtop homogeniser (Stretton Scientific Ltd) at 6500 rpm for 30 s. Following this, samples were passed through Qiashredders (Qiagen, Crawley, UK) and RNA was purified using the Qiagen RNeasy mini kit according to manufacturer’s instructions. Total RNA from exponentially growing WT or CHK1i resistant U20S was extracted using the PeqGold total RNA extraction kit (Peqlab), according to manufacturer’s instructions.

RNA was measured for purity and concentration with the NanoDrop1000 (ThermoFisher Scientific) and reverse transcribed using the Quantitect Reverse transcription Kit (Qiagen) according to manufacturer’s instructions. Quantitative real-time PCR was performed on 20ng cDNA, in triplicate, using predesigned Quanititect Primer assays (Qiagen) to the following murine genes; *Clspn, Chek1, Usp1, Usp14* and human genes *CLSPN, CHEK1, USP1* and *USP14.* These samples were run and analysed on a Rotor-gene Q system (Qiagen), using murine *Rpl13a* or human *RPL13A* primers as an internal control. All CT values were normalised to *Rpl13a*/*RPL13A* levels.

### Western blotting

Whole cell extracts were prepared from snap frozen pieces of Eµ-Myc, or Eµ-Myc*/c-Rel^-/-^* tumour tissue. Snap frozen tumour was lysed in PhosphoSafeM Extraction Reagent using the Precellys24 ceramic mix bead tubes (Stretton Scientific Ltd) in a Precellys®24 homogeniser (Stretton Scientific Ltd) at 6500 rpm for 30s, then extracted according to the PhosphoSafeM Extraction Reagent manufacturer’s instructions. In the case of cell lines samples, cell pellets were washed with ice-cold PBS, and lysed using PhosphoSafeM Extraction Reagent (Merck-Millipore, Watford, UK), according to manufacturer’s protocols. Protein quantification was undertaken using the BCA protein assay, and samples resolved by standard denaturing SDS-PAGE gels using the Criterion Gel System (Biorad). Samples were transferred onto PVDF membrane (Merck-Millipore) before being probed with the primary antibody. Horseradish peroxidase-conjugated secondary antibodies (anti-mouse; Sigma (UK), anti-rabbit; Sigma, UK) and enhanced chemiluminscence reagent (Thermo-scientific, UK) were used for detection.

### Antibodies

Antibodies used were CHK1 (phospho S345) (2341 Cell Signaling), CHK1 (2360 Cell Signaling), USP1 (14346-1-AP Proteintech), USP14 (14517-1-AP Proteintech), yH2AX (2577 Cell Signaling), GSK3B (phospho S9) (9336 Cell Signaling), GSK3B (9315 Cell Signaling), ERK 1/2 (phospho T202, Y204) (9101 Cell Signaling), ERK 1/2 (9102 Cell Signaling), AKT (phospho S473) (9271 Cell Signaling), AKT (phospho T308) (9275 Cell Signaling), AKT (9272 Cell Signaling), JNK (phospho T183, Y195) (9251 Cell Signaling), JNK (9252 Cell Signaling), p38 (phospho T180, Y182) (9211 Cell Signaling), p38 (9212 Cell Signaling), B-ACTlN (A5441 Sigma), CDC25B (9525 Cell Signaling), CHK2 (3440 Cell Signaling), PEA15 (2780 Cell Signaling), Pl3K (phospho Y458) (4228 Cell Signaling), Pl3K (4229 Cell Signaling), Karyopherin (sc-137016 Santa Cruz), CDK2 (2546 Cell Signaling), CDK1 (phospho Y15) (4539 Cell Signaling) and CDK1 (9116 Cell Signaling). Antibodies to the murine form of Claspin was generated by Moravian Biotechnologies. Anti-rabbit lgG (A6154 Sigma and 7074 Cell Signaling) and anti-mouse lgG (7076 Cell Signaling) HRP-linked secondary antibodies were used for western blot detection.

### Eµ-Myc mice studies

Eµ-Myc*/c-Rel^+/-^* offspring were generated by mating *c-Rel^-/-^* female mice with Eµ-Myc male mice, further Eµ-Myc*/c-Rel^-/-^* mice were generated by crossing Eµ-Myc*/c-Rel^+/-^* males with *c- Rel^-/-^* female mice. Eµ-Myc transgenic mice, and the associated crosses were monitored daily and were sacrificed at pre-determined end-points, defined as the animal becoming moribund, losing bodyweight/condition and/or having palpable tumour burden at any lymphoid organ site. Moribund mice were necropsied and single cell suspensions were prepared from tumour-bearing organs. Briefly, lymph nodes, spleen or thymus were homogenised through a cell strainer, and single cell suspension collected in DMEM (Lonza) supplemented with 10% FBS, 5mM L-glutamine, 5mM sodium pyruvate, 1μM L-asparagine and 50μM β-mercaptoethanol (Sigma Aldrich). These cell suspensions were then frozen in 90% FBS/10% DMSO for long- term storage and transplantation.

### Reimplantation studies

For tumour therapy studies, 2x10^6^ Eµ-Myc or Eµ-Myc*/c-Rel^-/-^* tumour cells from male mice were transplanted intravenously (IV) via the lateral tail vein into 8-week old male C57BL/6 recipients. Mice were monitored daily using parameters such as their bodyweight and food and water consumption to assess disease progression. Mice were necropsied when they became moribund and the tumour burden assessed.

Oral administration of the CHK1 inhibitor, CCT244747, prepared as previously described [25], or vehicle control (65% PEG-400, 20% Tween-20, 10% H_2_O, 5% DMSO (all Sigma Aldrich)) was initiated when tumours became palpable (approximately 10 days after inoculation of Eµ-Myc cells, and 20 days after inoculation of Eµ-Myc*/c-Rel^-/-^* cells). During efficacy studies, CCT244747 was given as a single agent, bolus dose (100 mg/kg p.o.) for 9 consecutive days. Lymphoid tumour burden and final tumour weights were measured at necropsy 24 hours after the final dose. For acute proteomic studies, CCT244747 was given as a single agent, bolus dose (100 mg/kg p.o.) once approximately 14 days after inoculation of Eµ-Myc cells and 25 days after inoculation of Eµ-Myc*/c-Rel^-/-^* cells, and the tumours necropsied either 8, 24 or 48 hours after dosing.

Oral administration of the PI3K inhibitor, GDC-0941, prepared as previously described [53], or vehicle control (0.5% methyl cellulose, 0.2 % Tween-80 (all Sigma Aldrich)) was initiated when tumours became palpable and given as a single agent, bolus dose (100 mg/kg p.o.) for 9 consecutive days. Lymphoid tumour burden and final tumour weights were measured at necropsy 24 hours after the final dose.

### Proteomics and analysis

Tissue extracts were prepared from snap frozen pieces of Eµ-Myc, or Eµ-Myc/c-Rel-/- splenic tumours. Briefly, tissue samples were suspended in 100 mM triethylammoniumbicarbonate (TEAB) with a mixture of protease and phosphatase inhibitors (cOmplete Mini EDTA-free protease inhibitor cocktail plus PhoSTOP phosphatase inhibitor cocktail, both obtained from Roche), homogenised by bead beater, and sonicated on ice. Lysed extracts were incubated with 0.1% (w/v) Rapigest SF (Waters) for 10 min at 80°C, left to cool, and incubated for 10 min on ice with Benzonase endonuclease (Merck Millipore) to digest nucleic acids. Samples were centrifuged (14,000*g*, 10 min at 4°C) to pellet cell debris. Protein concentration of the clarified lysate was ascertained by Bradford assay. Protein (200 µg) from each sample was aliquoted for protein digestion.

Oisulphide bonds were reduced (4 mM OTT in 100 mM TEAB, 10 min at 60°C) and free cysteines alkylated with iodoacetamide (14 mM in 100 mM TEAB, for 30 min, RT in the dark). lodoacetamide was quenched by addition of OTT to a final concentration of 7 mM. Proteins were digested with 2% (w/w) trypsin overnight at 37°C with gentle agitation. Resultant peptides were labelled with TMT 6-plex reagents (Thermo Scientific) at an 8:1 tag:protein ratio as per the manufacturer’s instructions, with labels assigned to samples randomly for the first biological replicate and shifted for each subsequent replicate. The labelling reaction was quenched by addition of 0.3% (v/v) hydroxylamine (Thermo Scientific) in 100 mM TEAB. TMT labelled peptides were mixed and dried to completion by vacuum centrifugation before re-suspending in 100 mM TEAB/ 1% TFA to hydrolyse the Rapigest SF (RT, 10 min). lnsoluble Rapigest SF cleavage product was removed by centrifugation (13,000*g* for 15 min at 4°C), and the sample desalted using C18 spin columns (Pierce, #89852) as per the manufacturers protocol, prior to strong cation exchange using stage tips (packed in-house with 5 disks per 200 µl tip as described previously [54] (Empore Supelco 47 mm Cation Exchange disk, #2251)). Each mixed labelled peptide sample was split across 8 tips, with peptides passed through the equilibrated stage tips twice. Bound peptides were eluted with 5% NH_4_OH (3 x 100 µl) and dried to completion using a vacuum centrifuge.

Peptides were fractionated using basic reverse-phase liquid chromatography as described [54], with 65 fractions collected, partially dried by vacuum centrifugation, and concatenated into 5 pools. For each pool, 5% was aliquoted and dried to completion prior to MS analysis. The remaining 95% was subjected to TiO_2_-based phosphopeptide enrichment, as described previously [55].

Total protein and phosphopeptide enriched fractions were analysed by LC-MS/MS using an UltiMate 3000 RSLCTM nano system (Dionex) coupled in-line with a Thermo Orbitrap Fusion Tribrid mass spectrometer (Thermo Scientific). Peptides were loaded onto the trapping column (PepMap100, C18, 300 µm x 5 mm, Thermo Scientific) using partial loop injection with 2% acetonitrile (ACN), 0.1% TFA at a flow rate of 9 µl/min for 7 minutes. Peptides were resolved on an analytical column (Easy-Spray C18, 75 µm x 500 mm, 2 µm bead diameter) using a gradient from 96.2% A (0.1% formic acid):3.8% B (80% ACN, 0.1% formic acid) to 50% B over either 120 minutes (single injection for phosphopeptide-enriched samples and two injections for total protein samples) or 240 minutes (single injection for total protein samples only) at a flow rate of 300 nl/min. Full MS1 spectra were acquired in the Orbitrap over *m/z* 375-2000 (60K resolution at m/z 200), with a maximum injection time of 50 ms and an ACG target of 4e5 ions. Data-dependent MS2 analysis was performed using a top speed approach (3 s cycle time) with peptides fragmented by collision-induced dissociation [56] at a normalised collision energy (NCE) of 35%, with fragment ions detected in the ion trap (maximum injection time of 50 ms, ACG target of 1e4). Following acquisition of each MS2 spectrum, a synchronous precursor selection (SPS) MS3 scan was performed on the top 10 most intense fragment ions, with SPS-MS3 precursors fragmented using higher energy collision-induced dissociation (HCD), at an NCE of 65%, and analysed using the Orbitrap over *m/z* 100-500 (50K resolution at m/z 200) with a maximum injection time of 105 ms and an ACG target of 1e5 [57, 58].

Analysis of MS data, with quantification of TMT reporter ion distributions, was performed using Proteome Discoverer 2.4 (PD 2.4) in conjunction with MASCOT (v2.6) and Percolator. For peptide identification from MS2 spectra, raw data files were converted to mzML format and searched in MASCOT against the Mouse UniProt reviewed database (Downloaded 25/04/2018; 16,966 sequences) with parameters set as follows: MS1 tolerance of 10 ppm; MS2 tolerance of 0.6 Da; enzyme specificity was set as trypsin with two missed cleavages allowed; carbamidomethylation of cysteine and TMT 6-plex modifications (on peptide N-termini and lysine side chains) were set as fixed modifications; oxidation of methionine and acetylation of protein N-termini were set as variable modifications, with the addition of phosphorylation (at serine, threonine or tyrosine residues) for phosphopeptide-enriched samples. Percolator was used for control of false discovery rates with a target FDR of 0.05. For phosphopeptide-enriched samples, the ptmRS node, operated in phosphoRS mode, was added to the PD 2.4 workflow for phosphosite localisation. In parallel with peptide identification, relative quantification of TMT 6-plex reporter ions was performed in PD 2.4 using the ’Reporter ions quantifier’ node, to quantify reporter ions from MS3 spectra with a peak integration tolerance of 20 ppm using the ’most confident centroid’ integration method.

Quantitative ratios were calculated for each biological replicate to look for protein/phosphopeptide changes upon Quantitative ratios were log2 transformed and, for all proteins/phosphopeptides quantified in at least 3 out of 5 bioreps, statistical analysis was performed in R using the LIMMA package, using a p ≤ 0.05 significance cut off. The mass spectrometry proteomics data have been deposited to the ProteomeXchange Consortium (http://proteomecentral.proteomexchange.org) via the PRIDE partner repository [59] with the dataset identifiers Project accession: PXD026203 & Project DOI: 10.6019/PXD026203. Access for reviewers can be obtained through: Username: Password: 5Ka35Kkf

### RNA-Seq and analysis

RNA was extracted as described above and sample quality analysed using Tapestation automated electrophoresis (Aglient) according to manufacturer’s instructions. Sample RNA Integrity Number (RIN) score exceeded 6 in all cases. RNA-Seq analysis was performed in the Illumina platform using TruSeq stranded mRNA-Seq at 25 million (75 bp) single reads per samples and sequencing performed on a NextSeq high-output 75 cycle run. The raw sequence data quality was first inspected using FastQC and MultiQC. Transcript counts were generated via Salmon [60] using Release M20 (GRCm38.p6) of the mouse genome (for the mouse samples) and Release 31 (GRCh38.p12) of the human genome (for the human samples).

The quantification files were imported into R for gene-level analyses using tximport [61] and the differential gene expression analyses were carried out using DESeq2 [62]. The data has been depositied on ENA (https://www.ebi.ac.uk/ena/submit/sra/#home) with the accession number PRJEB45284. Access for reviewers can be obtained through: Username: Webin-59077, Password: 2vTrAAF0a8

### STRING and Venn diagram analysis

STRING analysis was performed using version 11.0 at https://string-db.org/ [63]. Where indicated CHEK1 and AKT1 were manually added to the protein list to determine connections to phosphoproteins identified from the proteomics analysis. Details on proteins analysed and connections are in Supp Data File 2. Venn diagram analysis was performed at http://bioinformatics.psb.ugent.be/webtools/Venn/. For Venn diagrams examining phosphorylation, phosphosites identified from proteomics analysis rather than phosphopeptides were used as some phosphosites are associated with more than one phosphopeptide. Details on phosphosites analysed and are in Supp Data File 5.

### Statistical analysis

GraphPad Prism software (http://www.graphpad.com, V6.0) was used for statistical analysis. Except where stated in figure legends, unpaired t-tests or One-way ANOVA were used to calculate P values (P values of p<0.05 were considered significant).

## Supplementary Figures

**Supp Figure 1.**
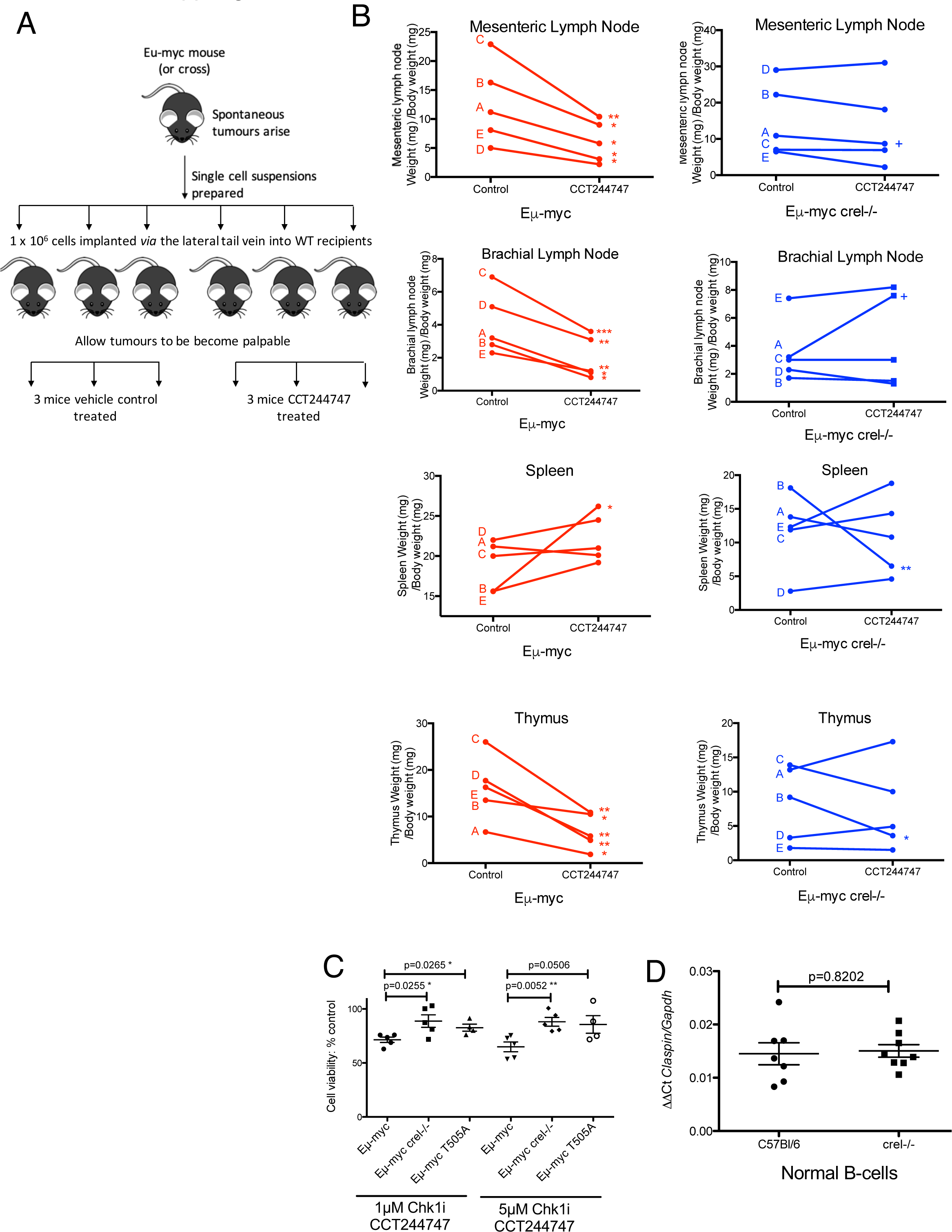
A) (B) Line graphs showing the mean response of the five reimplanted Eµ-Myc [64] and Eµ-Myc crel-/- (blue) tumours and their response to CCT244747 in further lymphoid organs. Each of the 5 tumours was implanted into 6 syngeneic recipient C57Bl/6 mice, 3 were treated with CCT244747 (100 mg/kg p.o), and 3 with vehicle control, for 9 days once tumours became palpable. A response was defined as a significant reduction (or increase) in tumour burden (P<0.05) using unpaired Student’s t-tests. (C) Eµ-Myc, Eµ-Myc/*c-re/-/- and* Eµ-Myc/*T505A* tumours were treated with 0.5 µM or 1 µM CCT244747 (or vehicle control) for 96 hours *ex vivo*. Eµ-Myc tumour cells show a reduced cell viability compared with the Eµ-Myc/*c-re/-/-* or Eµ-Myc/*T505A* tumour cells at all doses of CCT244747 tested. (D) Q-PCR showing relative Claspin expression in purified splenic B-cells from C57Bl/6 (n=6) and *cre/-/-* (n=7) mice. Data represents mean ± SEM.

**Supp Figure 2.**
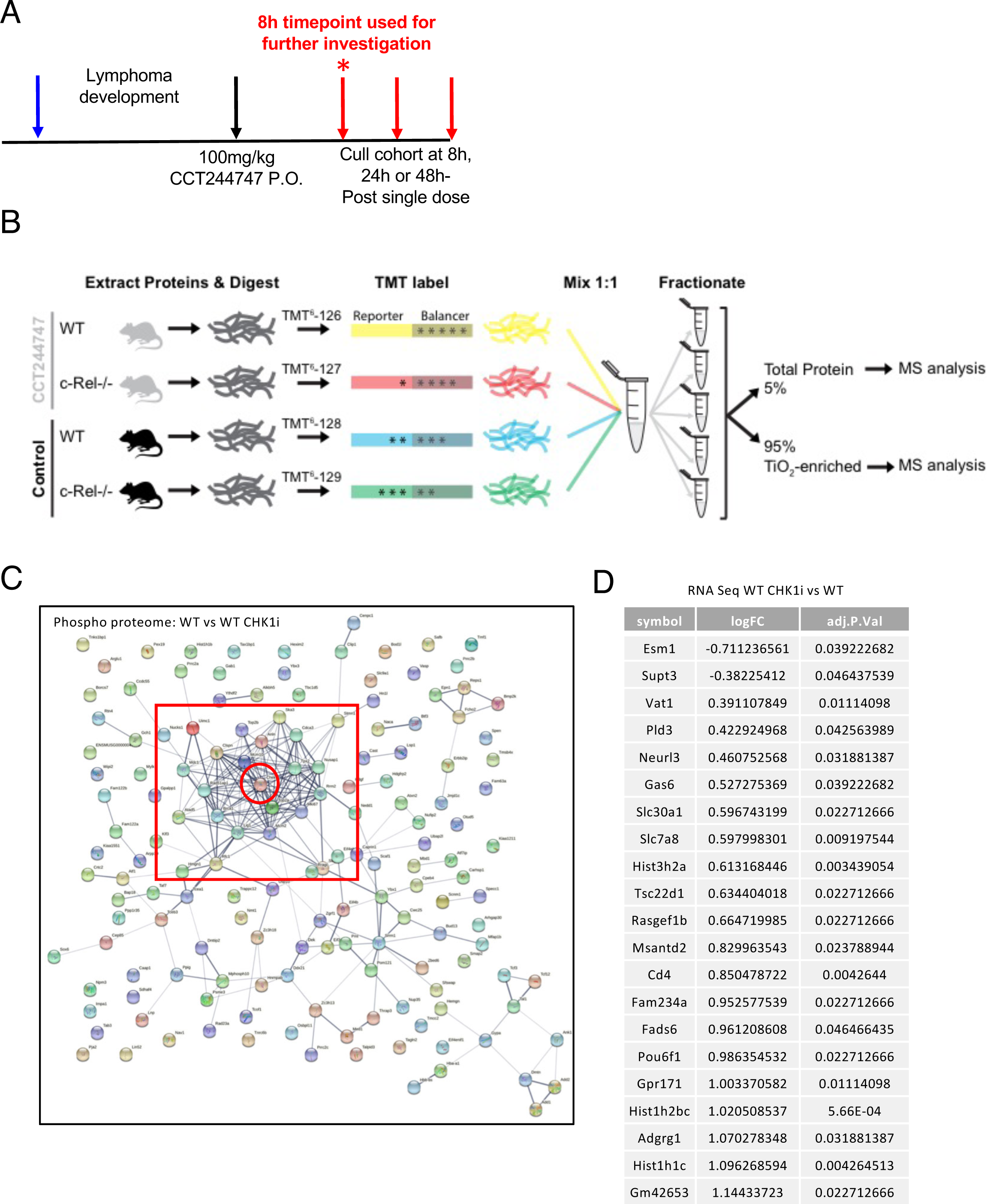
(A) Schematic diagram illustrating the single dose CHK1i *in vivo* study in Eµ-Myc and Eµ-Myc/*c-re/-/-* mice. 6 week old C57Bl/6 WT mice were implanted with either Eµ-Myc or Eµ-Myc/*c-re/-/-* (blue arrow) and once tumours became palpable were treated with either a single dose of 100 mg/kg CCT244747 p.o or vehicle control (black arrows), and then mice were then euthanised at either 8 hours, 24 hours or 48 hours later (red arrow). The star (*) denotes the 8 hour timepoint which was taken forward for use in future RNA-Seq and proteomic studies. (B) Schematic illustrating the workflow for proteomics experiments. Splenic tumours from Eµ-Myc or Eµ-Myc/c-Rel-/- mice were necropsied 8 hours post-treatment with either CHK1 inhibitor CCT244747 or vehicle control. Proteins were extracted and digested with trypsin prior to peptide labelling with tandem mass tags (TMT). Differentially labelled peptides from each treatment condition were mixed then fractionated via basic reverse-phase liquid chromatography, initially into 65 fractions which were concatenated into 5 pools. For each pool, 5% of the material was analysed by LC-MS/MS to obtain relative-quantification of total protein levels whilst the remaining 95% was subject to titanium dioxide (TiO_2_)-based phosphopeptide enrichment prior to LC-MS/MS analysis for phosphoproteomic analysis. (C) Wider STRING analysis showing that a cluster of proteins associated with CHK1 had down-regulated phosphorylation was upon CCT244747 treatment in Eµ-Myc WT tumours after a single dose of the CHK1i, as shown in Fig 2B. (D) Table detailing the 21 genes which showed differential expression in the Eµ-Myc WT tumours following acute CCT244747 treatment by RNA-Seq analysis.

**Supp Figure 3.**
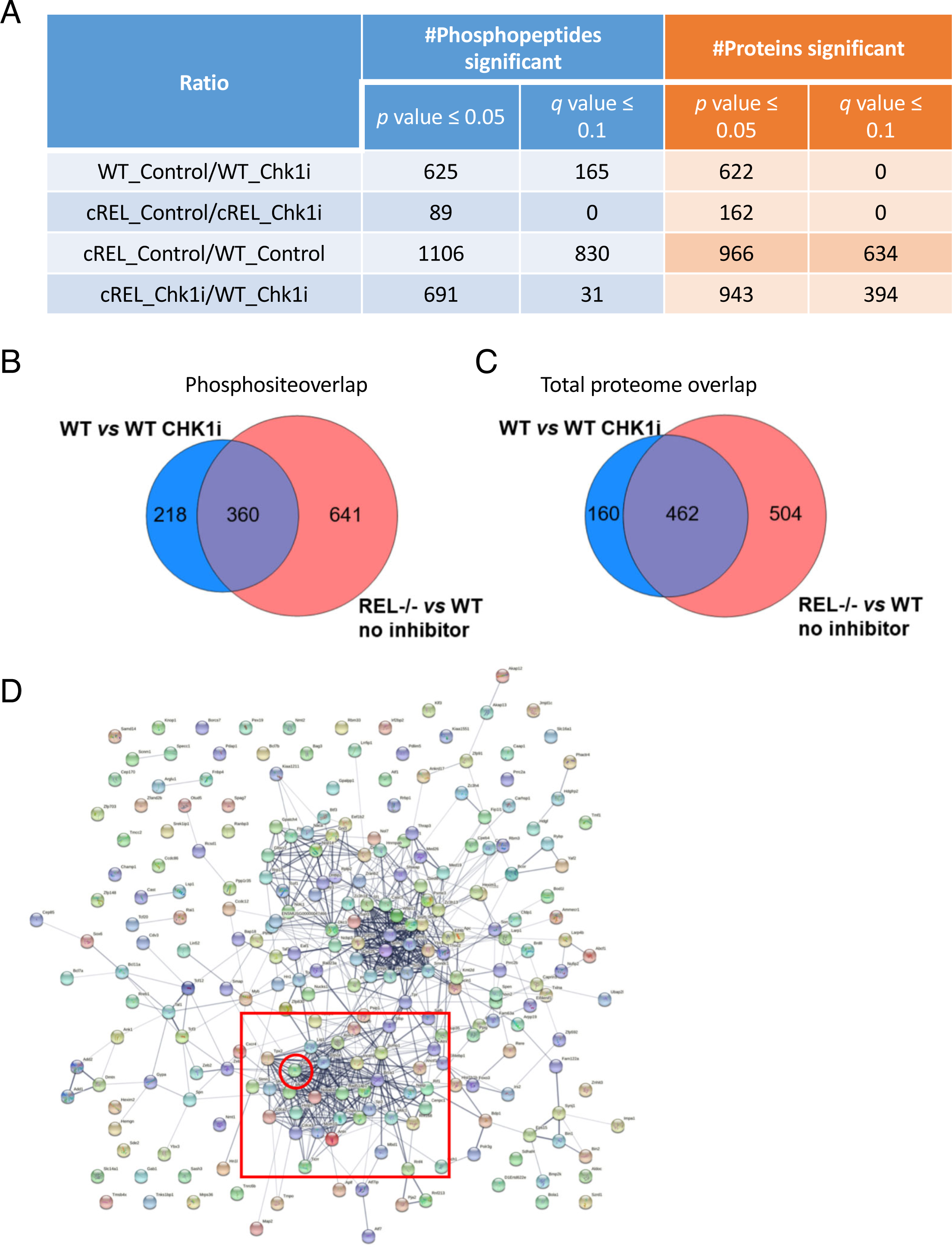
(A) Table detailing the number of phosphopeptides (orange column) and proteins (blue column) that were significantly different when the genotypes and/or treatment arms were compared after proteomic analysis in the acute CCT244747 study. A significant different was defined as an adjusted p value of <0.05 (B) Venn diagram illustrating that there is a high number of phosphosite changes in the c-Rel^-/-^ lymphomas without inhibitor treatment, but there is also a high level of overlap between these and the WT lymphomas following CCT244747 treatment. (C) Venn diagram illustrating that there is a high number of total protein changes in the c-Rel^-/-^ lymphomas without inhibitor treatment, but there is also a high level of overlap between these and the WT lymphomas following CCT244747 treatment. (D) Wider STRING analysis of the proteins associated with the 589 down-regulated phospho-peptides in the Eµ-Myc c-Rel^-/-^ lymphomas revealed that many of these had known linkages with CHK1 or CHK1 signalling, as shown in Fig 3D.

**Supp Figure 4.**
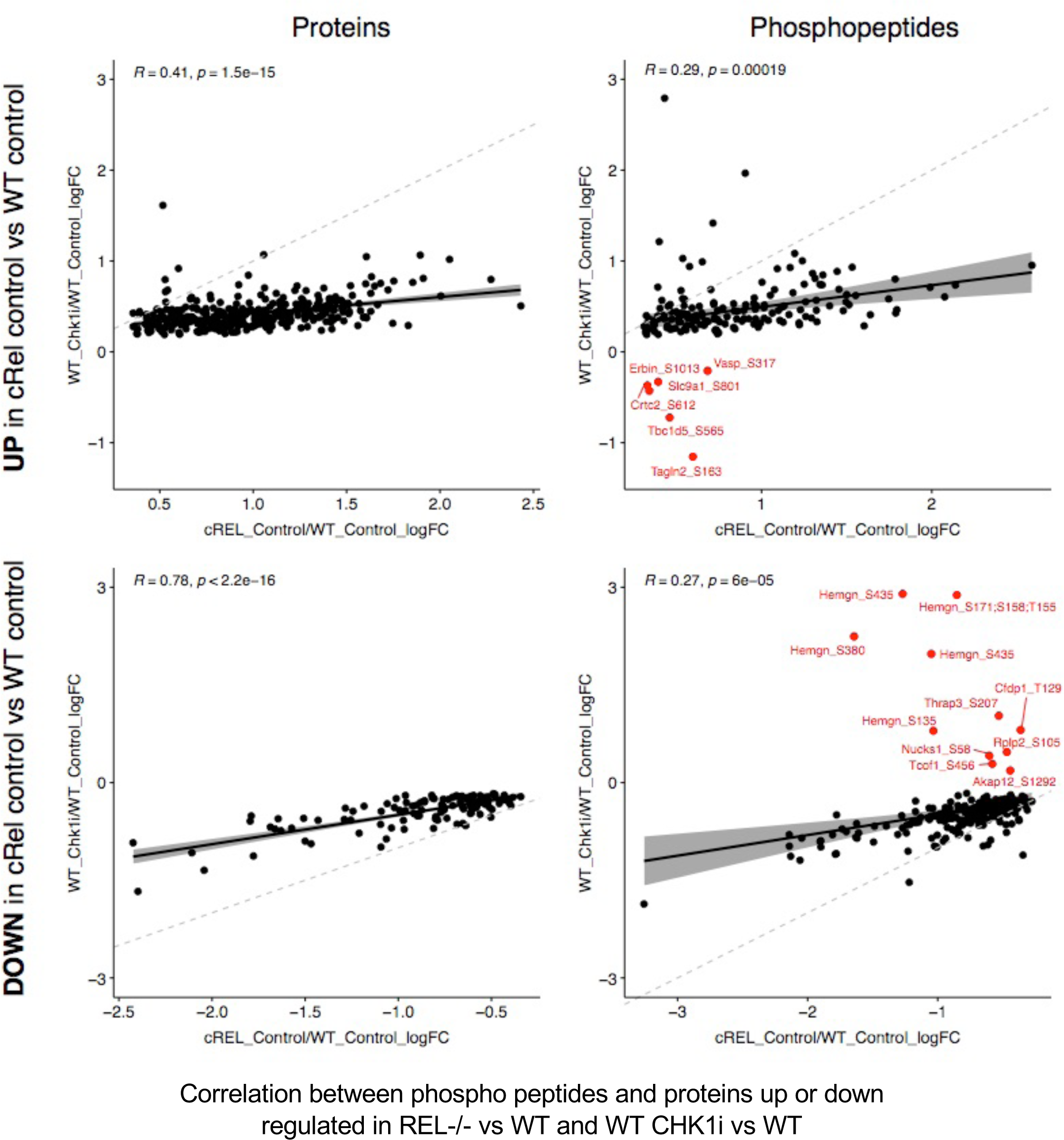
Correlation between phospho peptides and proteins up or down regulated in REL-/- vs WT and WT CHK1i vs WT

**Supp Figure 5.**
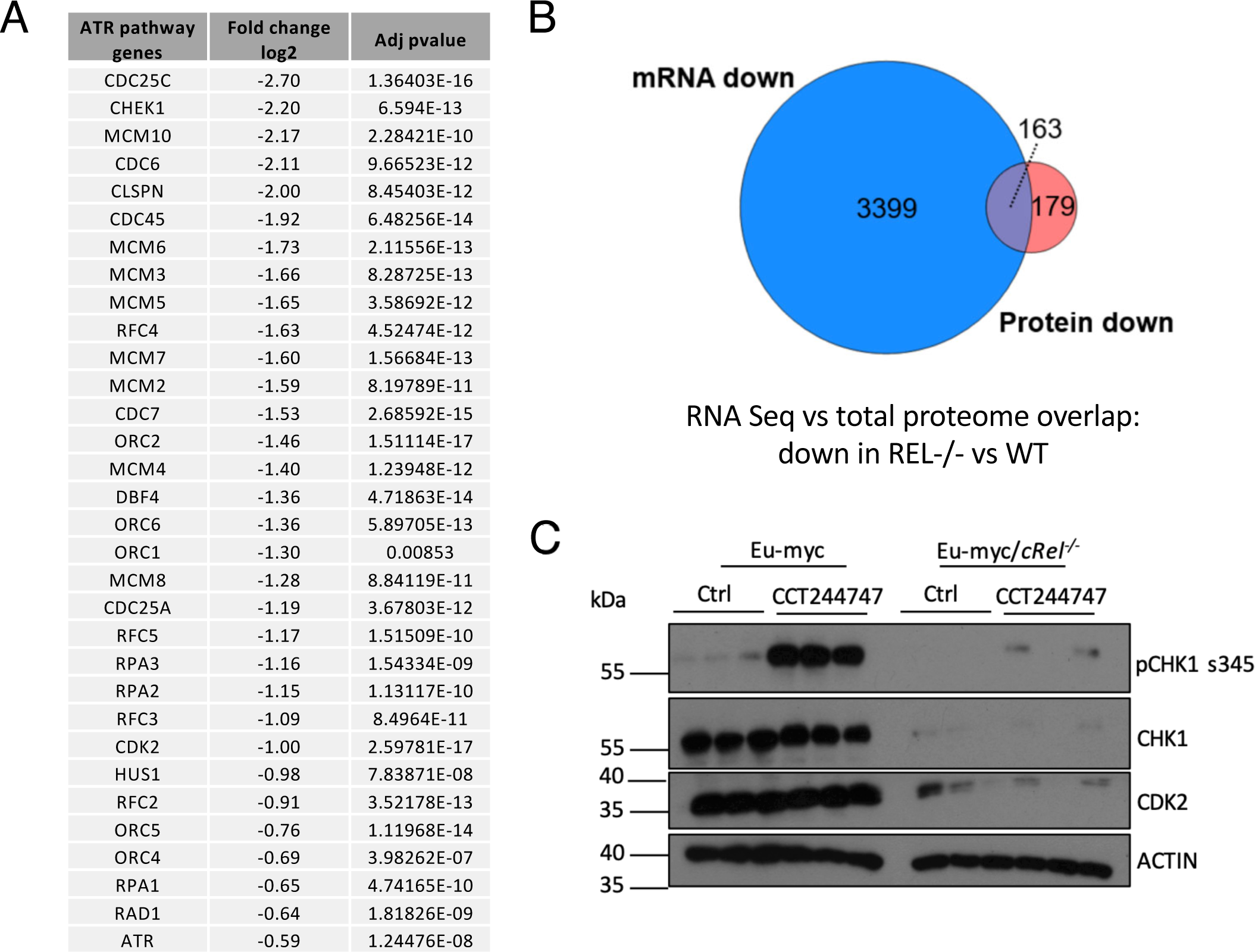
(A) Table showing the 32 genes associated with the ’Activation of ATR in response to replication stress’ that were downregulated at the transcript level in the Eµ-Myc c-Rel^-/-^ tumours when compared with Eµ-Myc WT tumours. A significant different was defined as an adjusted p value of <0.05 (B) Venn diagram showing the overlap between proteins down-regulated in the Eµ-Myc c-Rel^-/-^ tumours proteomic analysis and the RNA-Seq analysis, when compared with Eµ-Myc WT tumours. (C) Western blot analysis of CLASPIN, phospho-Ser345 CHK1, CHK1, CDK2 or ACTIN in snap frozen tumour extracts prepared from more re-implanted Eµ-Myc and Eµ-Myc/*c-re/-/-* tumours mouse inguinal lymph nodes 8 hours following a single dose of CCT244747. The expression of CHK1 and related pathway components are lost in Eµ-Myc/*c-re/-/-* tumours.

**Supp Figure 6.**
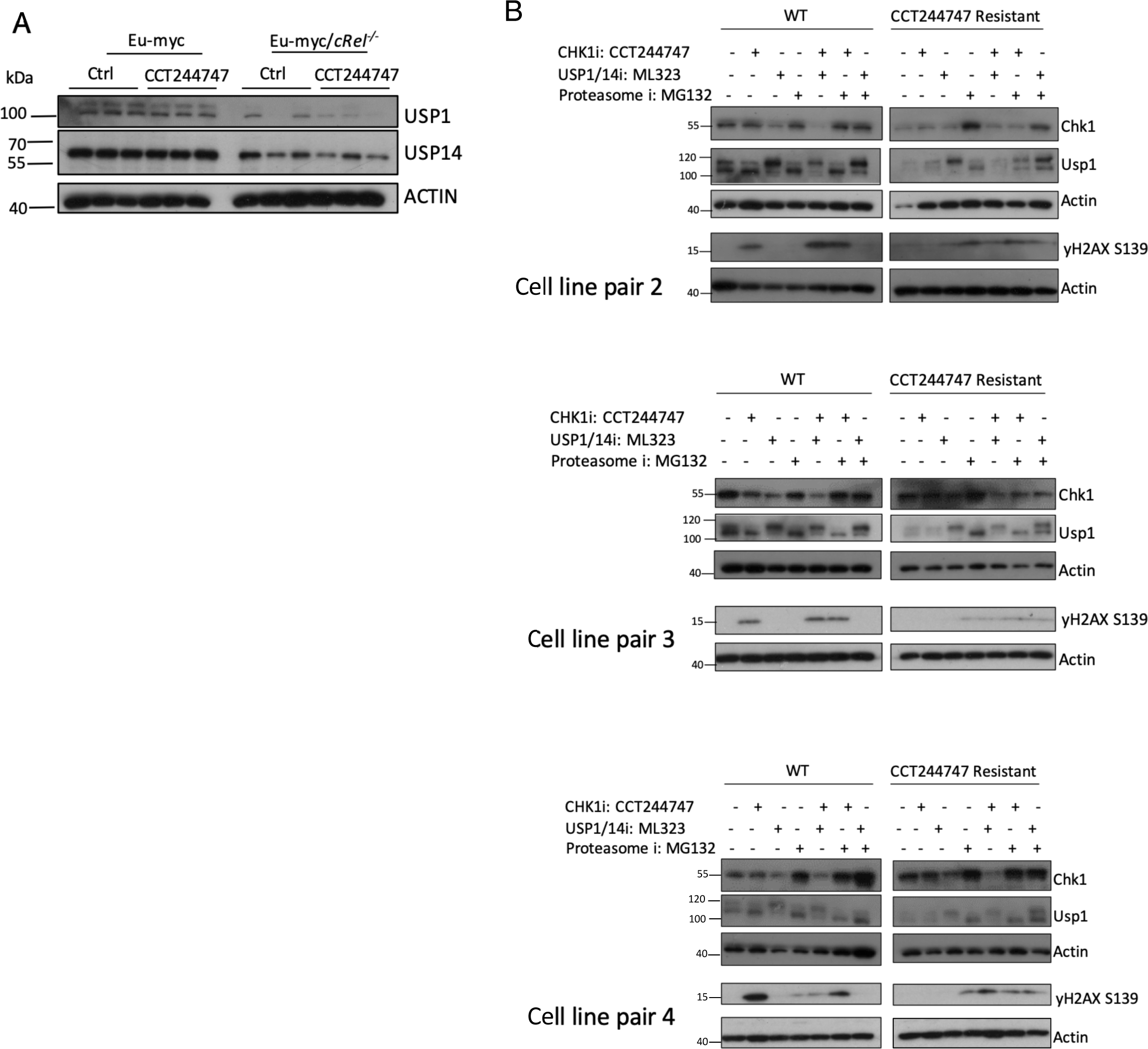
(A) Western blot analysis of USP1, USP14 or ACTIN in snap frozen tumour extracts prepared from more re-implanted Eµ-Myc and Eµ-Myc/*c-re/-/-* tumours mouse inguinal lymph nodes 8 hours following a single dose of CCT244747. USP1 and USP14 expression is lost in Eµ-Myc/*c-re/-/-* tumours. (B) Further western blot analysis of independent isolates of WT or CCT244747 resistant (CR) U20S cells treated with CCT244747, the USP1/14 inhibitor, ML323, or the Proteasome inhibitor MG-132, alone or in combination. Blots were probed for CHK1, USP1, yH2AX or ACTIN. Inhibition of USP1/14 in WT U20S results in the loss of CHK1. Proteasomal inhibition in the CCT244747 resistant U20S cells results in the stabilization of CHK1 protein.

**Supp Figure 7.**
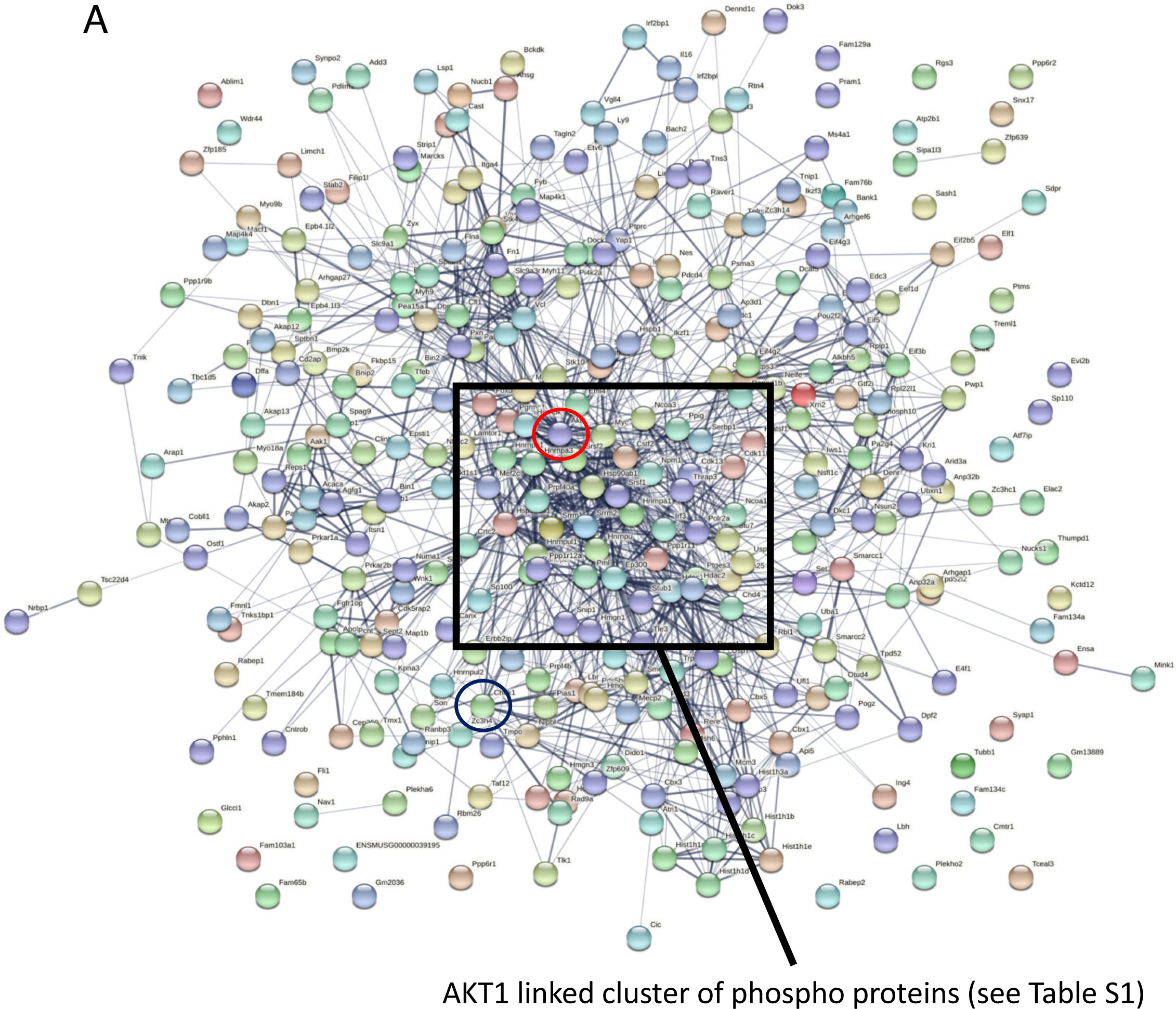
(A) STRING analysis showing that a cluster of proteins associated with AKT signalling is up-regulated was upon CCT244747 treatment in Eµ-Myc c-Rel^-/-^ tumours.

**Supp Figure 8.**
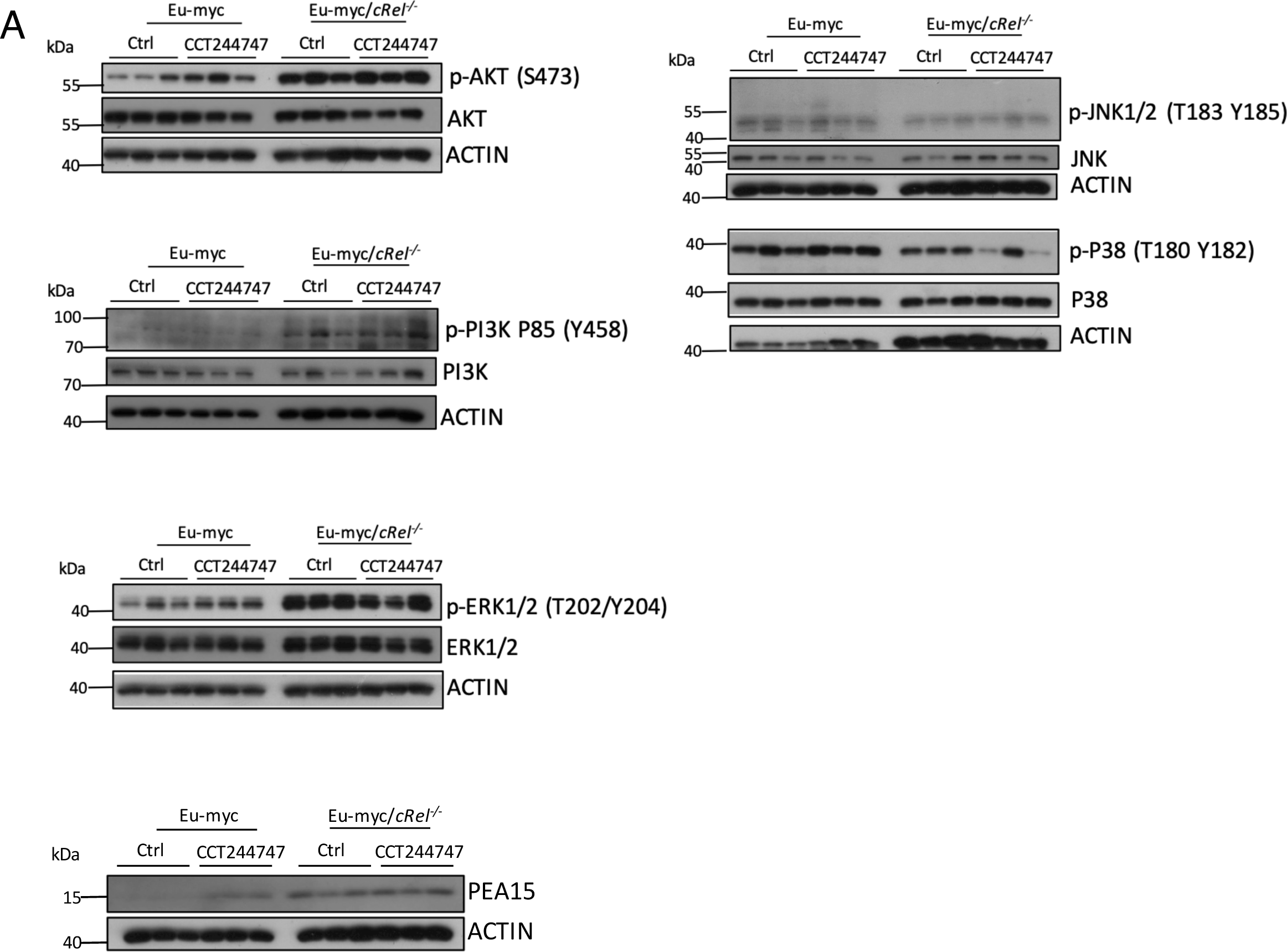
(A) Western blot analysis of phospho-Ser473 AKT, AKT, or ACTIN in snap frozen tumour extracts prepared from more re-implanted Eµ-Myc and Eµ-Myc/*c-re/-/-* tumours mouse inguinal lymph nodes 8 hours following a single dose of CCT244747. The data shows that the AKT pathway is highly in Eµ-Myc/*c-re/-/-* tumours. (B) Western blot analysis of phospho-Tyr458 PI3K, PI3K, or ACTIN in snap frozen tumour extracts prepared from re-implanted Eµ-Myc and Eµ-Myc/*c-re/-/-* tumours mouse inguinal lymph nodes 8 hours following a single dose of CCT244747. The data shows that the PI3K/AKT pathway is highly in Eµ-Myc/*c-re/-/-* tumours. (C) Western blot analysis of phospho-Thr202/Tyr204 ERK1/2, ERK1/2 or ACTIN in snap frozen tumour extracts prepared from more re-implanted Eµ-Myc and Eµ-Myc/*c-re/-/-* tumours mouse inguinal lymph nodes 8 hours following a single dose of CCT244747. The data shows that the ERK pathway is highly in Eµ-Myc/*c-re/-/-* tumours. (D) Western blot analysis of PEA15, or ACTIN in snap frozen tumour extracts prepared from re-implanted Eµ-Myc and Eµ-Myc/*c-re/-/-* tumours mouse inguinal lymph nodes 8 hours following a single dose of CCT244747. (E) Western blot analysis of phospho-Thr183/Tyr185 JNK1/2, JNK1/2, phospho-Thr180/Tyr182 p38, p38 or ACTIN in snap frozen tumour extracts prepared from re-implanted Eµ-Myc and Eµ-Myc/*c-re/-/-* tumours mouse inguinal lymph nodes 8 hours following a single dose of CCT244747. The data shows that the JNK/p38 pathway signalling is reduced in Eµ-Myc/*c-re/-/-* tumours.

**Supp Figure 9.**
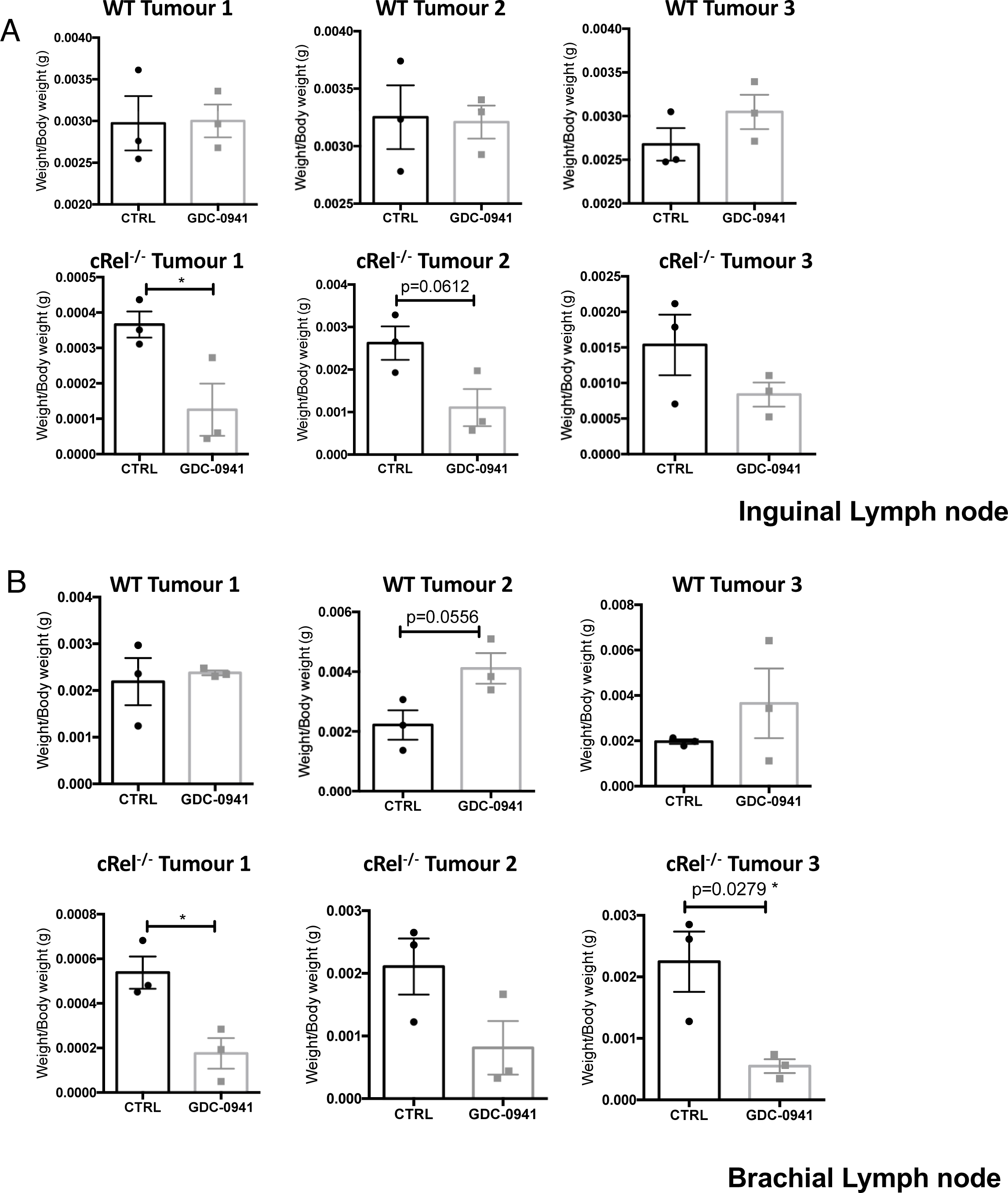
(A and B) Scatter showing the response of three reimplanted Eµ-Myc (top panel) and three Eµ-Myc cRel^-/-^ (bottom panel) tumours and their response to GDC-0941/Pictilisib in the inguinal lymph node (A) and brachial lymph node (B) tumour sites. Each of the 3 tumours was implanted into 6 syngeneic recipient C57Bl/6 mice, 3 were treated with GDC-0941/Pictilisib (100 mg/kg p.o), and 3 with vehicle control, for 9 days once tumours became palpable. A response was defined as a significant reduction (or increase) in tumour burden (P<0.05) using unpaired Student’s t-tests. WT Eµ-Myc showed little response to GDC-0941/Pictilisib whereas the Eµ-Myc cRel^-/-^ tumours were reduced by GDC-0941/Pictilisib.

**Supp Figure 10.**
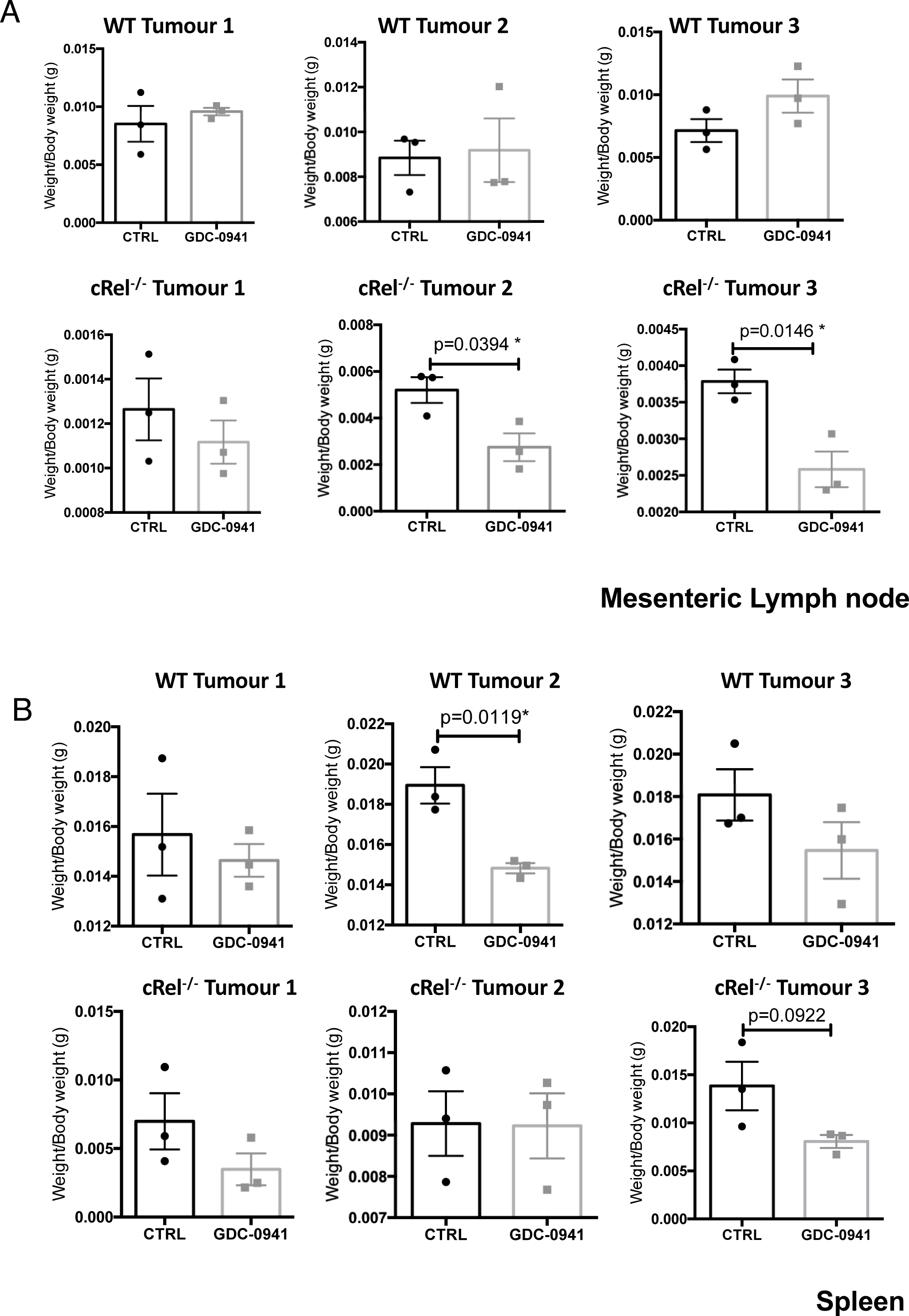
(A and B) Scatter showing the response of three reimplanted Eµ-Myc (top panel) and three Eµ-Myc cRel^-/-^ (bottom panel) tumours and their response to GDC-0941/Pictilisib in the mesenteric lymph node (A) and spleen (B) tumour sites. Each of the 3 tumours was implanted into 6 syngeneic recipient C57Bl/6 mice, 3 were treated with GDC-0941/Pictilisib (100 mg/kg p.o), and 3 with vehicle control, for 9 days once tumours became palpable. A response was defined as a significant reduction (or increase) in tumour burden (P<0.05) using unpaired Student’s t-tests. WT Eµ-Myc showed little response to GDC-0941/Pictilisib whereas the Eµ-Myc cRel^-/-^ tumours were reduced by GDC-0941/Pictilisib.

**Supp Table 1.**
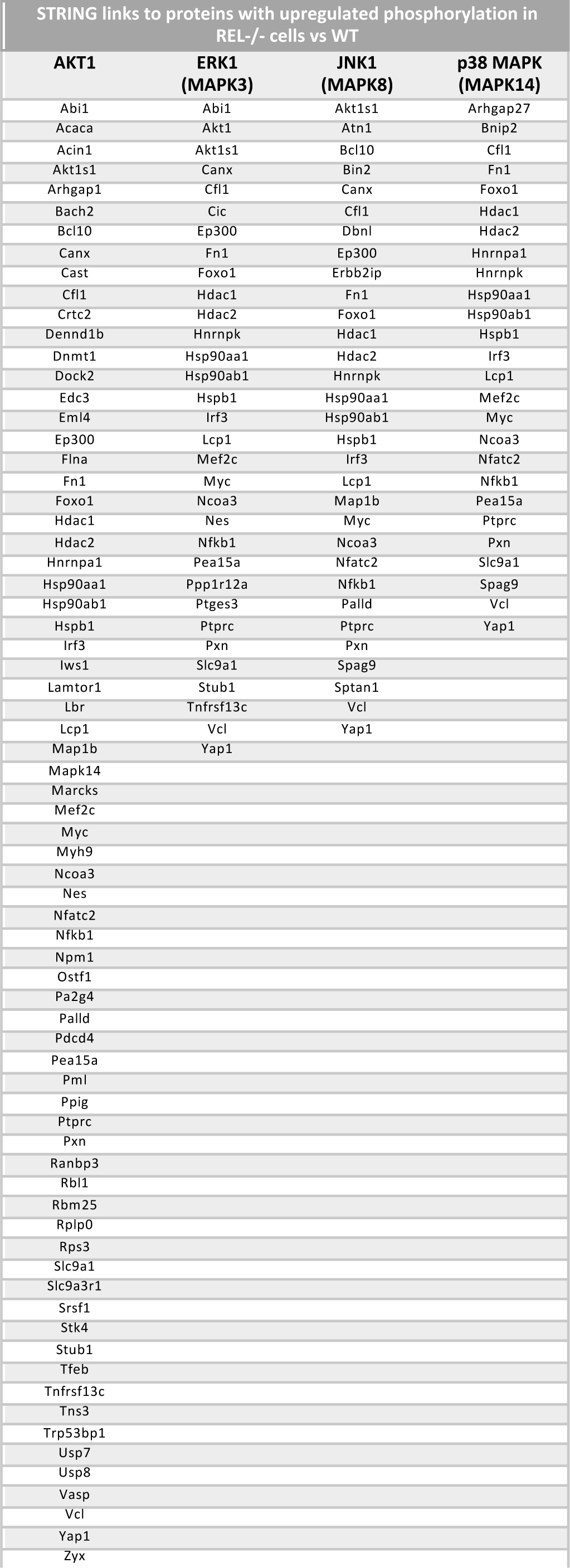
(A) Table detailing proteins in the AKT, ERK1, JNK and p38 pathways that had up-regulated phosphorylation when assessed by STRING analysis. The AKT1 pathway had the greatest number of links with a total of 71 proteins associated with AKT signaling showing increased phosphorylation

## Supplementary data files

Supp Data File 1 Proteomics.xlsx

Data from proteomics analysis of reimplanted Eμ-Myc lymphoma cells with either vehicle of CHK1i (CCT244747) treatment for 8 hours.

Tab1: Total protein data

Tab 2: Phosphopeptide data

Supp Data File 2 STRING interactions.xlsx

STRING interactions data based phosphorylated proteins identified from phospho proteomics analysis.

Tab 1: Analysis of STRING interactions between phospho proteins down regulated in WT Eμ-Myc lymphoma cells after CHK1i (CCT244747) treatment. Note CHK1 (Chek1) was manually added into the protein list in order to identify potential interactions with the target of the CCT244747.

Tab 2: Analysis of STRING interactions between phospho proteins down regulated in c-Rel null versus WT Eμ-Myc lymphomas. Note CHK1 (Chek1) was manually added into the protein list in order to identify potential interactions with the target of the CCT244747.

Tab 3: Analysis of STRING interactions between phospho proteins up regulated in c-Rel null versus WT Eμ-Myc lymphomas. Note AKT1 was manually added into the protein list in order to identify potential interactions with this kinase.

Supp Data File 3 RNASeq_all_genes_list_EuMyc.xlsx

Gene lists from RNA Seq analysis of reimplanted Eμ-Myc lymphoma cells with either vehicle of CHK1i (CCT244747) treatment for 8 hours.

Tab 1: Comparison of gene expression changes in WT Eμ-Myc lymphoma cells with either vehicle or after after CHK1i (CCT244747) treatment

Tab 2: Comparison of gene expression changes between WT and c-Rel null Eμ-Myc lymphoma cells after CHK1i (CCT244747) treatment

Tab 3: Comparison of gene expression changes between WT and c-Rel null Eμ-Myc lymphoma cells with no CHK1i (CCT244747) treatment

Supp Data File 4 RNASeq_counts_tximport_EuMyc.csv

Data for all genes and samples from RNA Seq analysis of reimplanted Eμ-Myc lymphoma cells

Supp Data File 5 Venn diagrams.xlsx

Data files from Venn analysis of Eμ-Myc lymphoma cell proteomics and RNA Seq data

Tab1: Comparison between phosphosites downregulated in untreated c-Rel null relative to untreated wild type with those down regulated in WT cells treated with CHK1i (CCT244747) versus untreated WT.

Tab 2: Comparison between total proteins downregulated in untreated c-Rel null relative to untreated wild type with those down regulated in WT cells treated with CHK1i (CCT244747) versus untreated WT.

Tab3: Comparison between all phosphosite changed in untreated c-Rel null relative to untreated wild type with those down regulated in WT cells treated with CHK1i (CCT244747) versus untreated WT.

Tab 4: Comparison between all total changesin untreated c-Rel null relative to untreated wild type with those down regulated in WT cells treated with CHK1i (CCT244747) versus untreated WT.

Tab 5: Comparison between mRNAs and proteins down regulated in untreated c-Rel null relative to untreated wild type

Tab6: Comparison between phosphosites upregulated in untreated c-Rel null relative to untreated wild type with those down regulated in WT cells treated with CHK1i (CCT244747) versus untreated WT.

Tab 7: Comparison between total proteins upregulated in untreated c-Rel null relative to untreated wild type with those down regulated in WT cells treated with CHK1i (CCT244747) versus untreated WT.

Supp Data File 6 RNASeq all_genes_list_U20S.xlsx

Gene lists from RNA Seq analysis of wild type control and CHK1i (CCT244747) resistant U20S cells with or without CCT244747 treatment for 24 hours.

Tab 1: Comparison of gene expression changes in between WT untreated U20S cells (WT.Not) and untreated CHK1i resistant U20S cells (CR.Not)

Tab 2: Comparison of gene expression changes in between WT untreated U20S cells (WT.Not) and WT CHK1i treated U20S cells (WT.CHK1i)

Tab 3: Comparison of gene expression changes in between untreated CHK1i resistant U20S cells (CR.Not) and CHK1i treated resistant U20S cells U20S cells (CR.CHK1i)

Tab 4: Comparison of gene expression changes in between CHK1i treated WT U20S cells (WT.CHK1i) and CHK1i treated CHK1i resistant U20S cells (CR.CHK1i)

Tab 5: Comparison of gene expression changes in between untreated WT U20S cells (WT.Not) and CHK1i treated CHK1i resistant U20S cells (CR.CHK1i)

Tab 6: Comparison of gene expression changes in between CHK1i treated WT U20S cells (WT.CHK1i) and untreated CHK1i resistant U20S cells (CR.CHK1i)

Supp Data File 7 RNA Seq counts_tximport_U20S.csv

Data for all genes and samples from RNA Seq analysis of U20S cells.

